# Structure learning and context inference emerge from next-state prediction in a cerebellar-medial temporal lobe circuit

**DOI:** 10.64898/2026.11.14.765964

**Authors:** Cheshta Bhatia, Samuel J. Gershman, Elom A. Amematsro

## Abstract

Humans learn many context-dependent behaviors; yet the contexts driving these behavior are rarely explicit and must instead be inferred. Further, we are rarely exposed to the same context multiple times, so we must learn how to generalize to new contexts from our experiences in prior ones. Motor learning relies on both. People can hold multiple conflicting visuomotor mappings and recall the appropriate mapping when needed. Much as cricket is easier to learn for someone who already plays baseball, experience with previous mappings accelerates learning of others. Indeed, human motor learning is well described by context inference, in which the learner infers the context and uses it to update and recall the appropriate memory. How the brain implements this inference, and which circuits perform it, however, is unknown. Here we introduce a circuit in which a cerebellar forward model predicts the next state conditioned on a latent from the medial temporal lobe. At the same time, the medial temporal lobe updates that latent from the cerebellum’s prediction error. While we only trained the circuit to predict next state, it learns to infer context and represents contexts in a structured latent space. Coupled with this latent, a motor cortex module reproduces context-dependent retrieval, savings after washout, spontaneous recovery, and evoked recovery, and compensates for perturbations it has never experienced. The same circuit reproduces cerebellar-dependence of object-location memory and independence in object-recognition memory, loss of place-cell stability in the dark when cerebellar plasticity is blocked, and the abstract geometry of hippocampal activity in a reversal task. Cerebellar–medial temporal lobe interaction is therefore a candidate locus for context inference and structure learning in the brain.

## 1 Introduction

Human behavior is context-dependent. For example, despite being in the same physical environment, individuals behave differently at a restaurant depending on whether they are dining with high school friends, coworkers, or their supervisor. While this context-dependence allows for flexible behavior, it also presents significant challenges. First, context is rarely explicitly cued and must instead be inferred from signals in the environment. Second, because we rarely encounter the same context multiple times, most of our experiences involve novel contexts. Thus the brain must learn how to act in these novel circumstances by generalizing from previous experiences: a process known as structure learning. Both processes, context inference and structure learning, are present in motor learning ^1,2^. However, how and where these computations are performed in the brain remain unclear. Given the importance of context inference and structure learning to motor and non-motor tasks, might a shared circuit support both? If so, which brain regions comprise this circuit? We hypothesized that a cerebellar-medial temporal lobe circuit supports context inference and structure learning in the brain.

Both brain regions have been implicated in context-dependent behavior and are known to interact. The cerebellum contains multiple internal models and switches between them based on context ^3^,^4^. Hippocampal activity remaps across contexts ^5^–^7^, represents context abstractly ^8^, and hippocampal lesions impair contextspecific memory ^9,10^. Further, the two regions are reciprocally connected ^11,12^, and cerebellar perturbation alters hippocampal activity ^13,14^. Thus there is significant evidence for the existence of the cerebellar-medial temporal lobe circuit. We thus asked how and if this circuit could support context inference.

Imagine helping a friend move a series of boxes: each containing an unknown mixture of books, clothes, and plates (Fig. 1a). The contents of each box are unknown but determine how we should carry each. For instance, some boxes are light while others may be heavy or lopsided. While we may not know the contents of the box before we pick it up, we can infer its contents once we do. We can formalize this intuition through the notion of forward models ^15,16^, which predict the next observation of an environment given the current observation and action taken by the agent. In the moving boxes example, the next observation could be the height of the box after we lift it, and the action could be how hard we lift it. Once the next observation is made, we can update our belief about the contents of the box from the difference between our prediction and the actual outcome (Fig. 1b). When the forward model is Gaussian, this update has three properties that are useful for context inference circuit. First, its only new external input is the prediction error of the observation. Its remaining inputs are the previous observation and action. Second, the update is Markovian and depends only on the previous state of *z* and the inputs. Finally, the latent that encodes context is continuous and can thus represent infinitely many contexts as well as the relations between them.

Each component of this latent inference circuit has a direct biological analog. The cerebellum is a forward model whose output tracks prediction error ^17–19^, and it connects to the medial temporal lobe ^11^, which is recurrent ^20^ and encodes context during inference tasks ^8^. While the function of cerebellar-medial temporal lobe interactions is unclear, the circuit they form has the structure of the latent-inference circuit above. Thus we hypothesized that cerebellar-medial temporal lobe interactions could mediate context inference and structure learning across motor control and cognition. To test this, we built a circuit from known components of the cerebellar-medial temporal lobe anatomy and asked whether this circuit was sufficient for context inference and structure learning over a set of motor tasks.

Over twelve unique motor contexts, spanning visuomotor rotations, velocity-dependent curl fields, and drifts, we found that MTL learned to extract the underlying context of each task. Further, MTL represented each context in a structured manner such that contexts from distinct perturbation families clustered in their own subspaces. Within each family subspace contexts were additionally arranged relationally, according to their parameters. We found that a motor cortex module receiving this input could reproduce context-dependent features of motor learning such as context-dependent retrieval, savings after long washout, spontaneous recovery, and evoked recovery. The structured representations also enabled motor cortex to generalize, compensating for novel contexts on the first trial. This generalization depended on the structure of pre-training, as in human structural learning ^1^. We then tested the same circuit on three memory tasks, object-location memory, object-recognition memory ^14^, and place-field stability in the dark ^13^, and found that it reproduced the pattern of cerebellar dependence in each. The circuit also reproduced hippocampal geometry during a cognitive context inference task ^8^. In each case, the medial temporal lobe extracted a latent context from the prediction errors of the cerebellum, which in turn depended on the outputs of the medial temporal lobe to learn the context-dependent consequence of actions. The circuit therefore provides a circuit-level mechanism by which the brain can perform context inference and structural learning, and a unified hypothesis for the function of cerebellar-medial temporal lobe interactions across disparate tasks.

**Figure 1.**
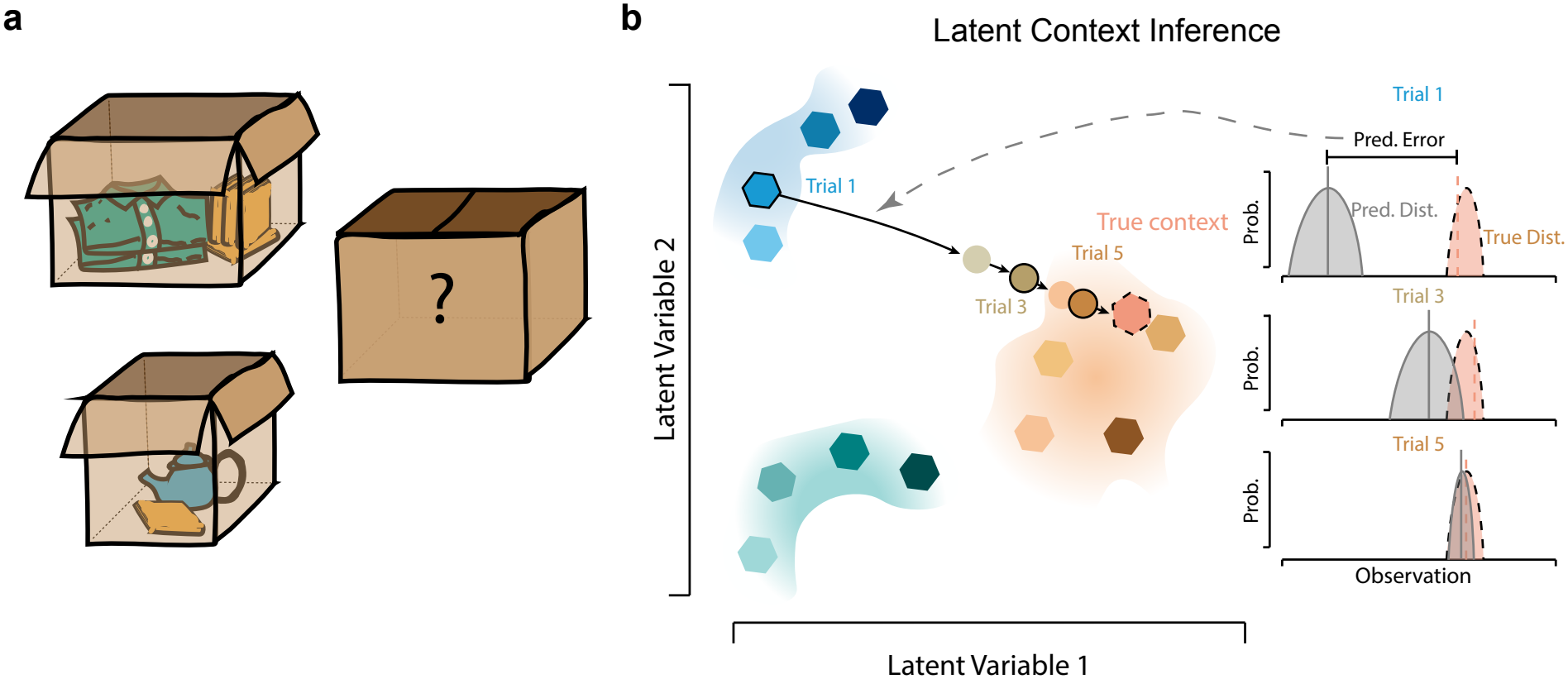
Latent Inference Through Prediction Errors. **a**, The contents of each box are hidden and fixed, but they determine what happens when the box is lifted. **b**, Latent context inference: the agent’s belief over latent variables is updated from the prediction error at each trial.

## 2 Results

### 2.1 A cerebellar–MTL circuit

We ask whether a circuit built from established components, trained to predict its next observation, is sufficient for context inference. The circuit has three modules (Fig. 2a). The cerebellar module (CB) is a two-layer feedforward network that receives the current observation (*o*_*t*_), an efference copy of the action (*a*_*t*_), and a latent signal (*z*_*t*_) from the medial temporal lobe module (MTL) and predicts the next observation (*o*_*t*+1_). The MTL module is a Gated Recurrent Unit that receives the previous observation and action (*o*_*t*−1_, *a*_*t*−1_), together with the cerebellum’s current prediction error *e*_*t*_, and outputs a 32-dimensional latent *z*_*t*_. CB and MTL form a loop; CB’s prediction error serves as MTL’s inputs, and MTL’s output serves as one of CB’s inputs. Both are trained only to predict the next observation. The third module, motor cortex (MC), is a feedforward policy with a single hidden layer that receives the current observation, the MTL output, and a 32-dimensional projection of the hidden state of CB. MC is trained via reinforcement learning to steer the agent towards randomly placed goals (Methods).

To isolate what each module contributes, we compare four circuit variants. The first circuit variant, Motor Cortex (MC) is the policy alone. It can select actions but has no forward model and no way to extract a context signal. Motor Cortex + Cerebellum (MC + CB) adds a forward model but no latent. Because a frozen cerebellum without a latent cannot adapt to a new context, CB is trained alongside MC in this circuit. COIN has a forward model and context inference implemented as a posterior over the set of trained contexts. It is given both the ground-truth forward model and the identities of the trained contexts, so it acts as an ideal observer with respect to context. Our implementation of COIN differs from the original implementation of Heald et al. ^2^ in two ways. First, we use a finite hypothesis set rather than the unbounded Dirichlet process. We found that the finite hypothesis better inferred novel contexts; however, none of our results depended on this. Second, rather than maintaining a set of independent policies, we use COIN as an input to a single context-conditioned policy. This means that all of our policies are implemented by a single module, MC, and thus different contexts can interfere with each other through these shared weights. This choice was motivated by the observation of context-dependent control in motor cortex ^21,22^, and we think is more biological. The final circuit, CB-MTL is the full circuit shown in Figure 2. It has a forward model and a continuous latent that may or may not support context inference. Comparing MC to MC + CB isolates the forward model; MC + CB to CB–MTL isolates the recurrent latent; CB–MTL to COIN contrasts a learned continuous latent with a normative posterior over a discrete set.

### 2.2 Prediction task

We hypothesized that training the CB-MTL circuit on next-state prediction—in a task where the future observation depended on a hidden context—would be sufficient to drive context inference in MTL’s latent, and because that latent is continuous it should also represent the relations amongst the contexts. We test the first claim here and the second in the following section. We train CB and MTL on next-state prediction across twelve uncued contexts and ask whether the CB-MTL circuit predicts better than an ideal context-free model, and whether that advantage depends on MTL.

The agent moves through a two-dimensional arena, under random actions drawn from a Gaussian process. The affect of these actions depend on a hidden context *k* that switches with probability 0.1 on each step. In simplified form, the dynamics are

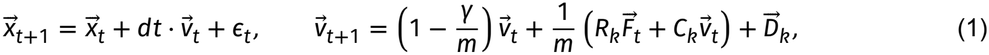

where 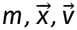, and 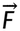 are the mass, position, velocity, and actions of the agent and *ϵ*_*t*_ is signal-dependent motor noise. The context sets {*R*_*k*_, *C*_*k*_, *D*_*k*_}, which define three families of perturbations (Extended Data Fig. 1): rotations, curls, and drifts respectively. We trained the model on four contexts from each family—rotations of ±30°, ±60°, curls of *κ* = ±7, ±13, and drifts of 0°, 90°, 180°, and 270°, and CB predicts the next position and velocity from the current position and velocity, action, and latent.

For the CB-MTL circuit, held-out velocity error fell from 0.0348 ± 0.0006 to 0.0138 ± 0.003 over training (Fig. 2b). A context-free model, which averages the twelve context-conditioned predictions, reaches 0.0223 ± 0.00049: similar to the error reached by a model without MTL (0.0228 ± 0.0002; Fig. 2b, purple). Thus next-state prediction alone is sufficient to drive the circuit to extract and leverage context information, and MTL is necessary for it.

Context was represented in the MTL latent, *z*. Projecting activity onto its leading principal components revealed twelve distinct clusters of activity: each corresponding to a unique context (Extended Data Fig. 2a). Context accounted for >50% of the variance of all of its principal components (Extended Data Fig. 2d). In contrast, the MTL hidden state reflected heading direction and speed, and the cerebellum represented position (Extended Data Fig. 2b,c,e,f.)

Accordingly, context could be linearly decoded from *z*. On held-out data, linear decoders were 60.6 ± 2.4% (S.D) accurate (chance 8.3%; Fig. 2e). Decoding errors were structured based on family and sign within a family. Across all twelve contexts, 39.3 ± 0.9% of errors stayed within the true family (chance 27.3%); within family, 98.1 ±0.3% of errors were magnitude errors that preserve the sign of the perturbation (chance 33.3%;). A −60° rotation is confused with a −30° rotation more often than with a 30°.

Decoder accuracy tracked the circuit’s ability to predict next-state. Both increased over training (Fig. 2c), and similarly depended on the prediction error and MTL. Ablating the prediction error of a trained model decreased accuracy by 36.7 ± 3.7% (relative; S.D) while increasing the prediction error by 44.3 ± 1.8% (Fig. 2d,e). Similarly, adding transient noise to MTL units decreased decoding accuracy by 45.8 ± 3.1% and increased prediction error by 24.9 ± 6.1%. Shuffling the prediction error of one trial with the prediction error of a different trial had the strongest effect on both: increasing the prediction error above the context-free floor (Fig. 2d). Decoding accuracy decreased by 65.7 ± 2.2%, and the prediction error increased by 78.2 ± 2.7%. Thus both the CB-provided prediction error and MTL integration causally contribute to extracting context and predicting the next state.

The CB-MTL circuit therefore learns to extract and represent each of the twelve trained contexts. The next question is whether it can represent novel contexts in a similarly structured manner.

**Figure 2.**
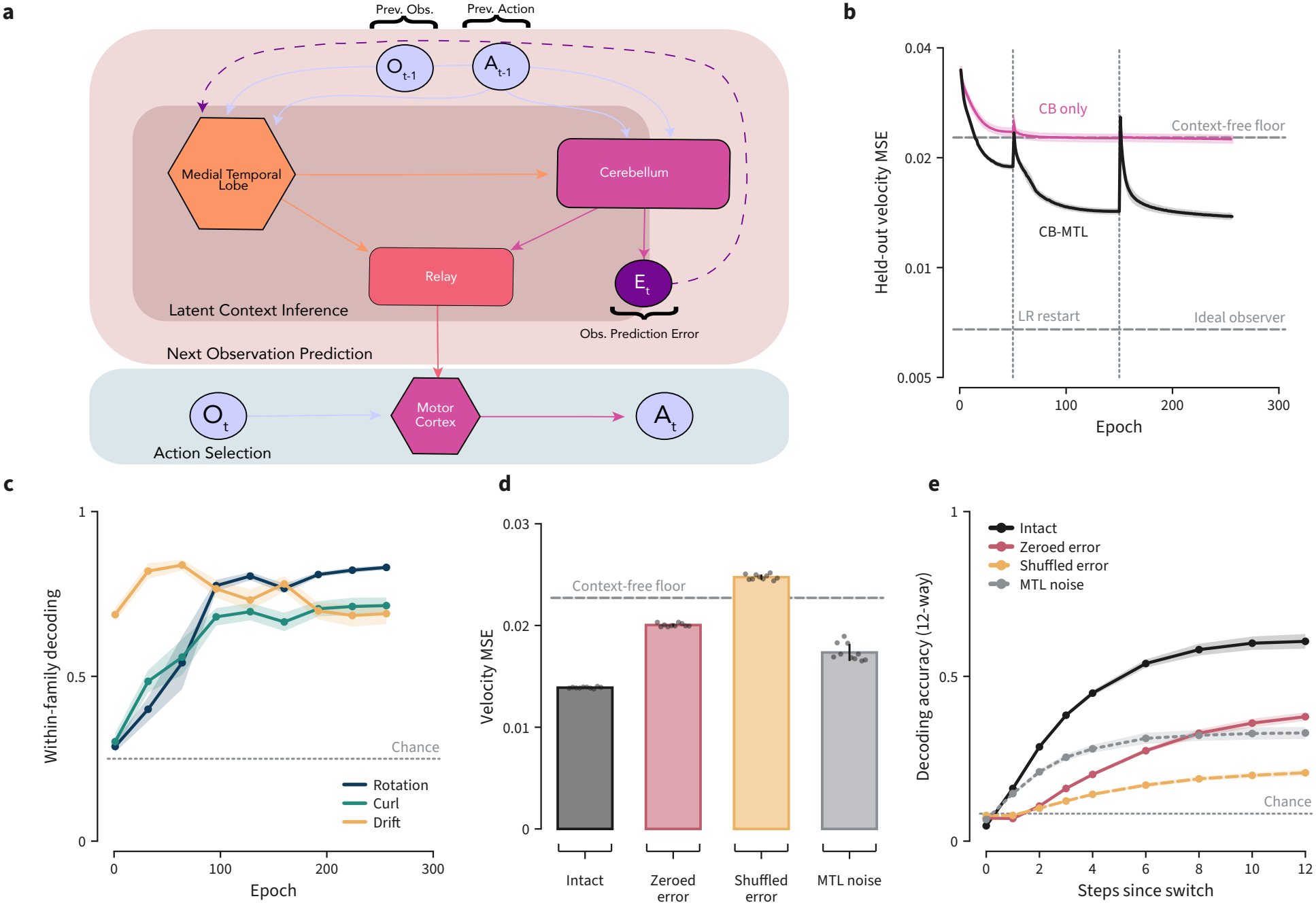
a, Circuit schematic. CB predicts the next observation from state, efference copy and the MTL latent; its prediction error is MTL’s input; both modules are trained only to reduce that error and are frozen thereafter. **b**, Held-out velocity prediction error over training, overall and separately for each family. Error falls from 0.0348 ± 0.0006 (S.D) to 0.0138 ± 0.003 over 256 epochs; a context-free network reaches 0.02278 ± 0.00015, close to the theoretical context-free floor of 0.02230 ± 0.00049. **c**, Within-family decoding of context from the MTL output over the same period; dotted line, chance. **d**, Mean squared error of the next-step velocity prediction for the same conditions; dots are individual networks. The prediction quality degrades when the error supplied from the cerebellum to the MTL is shuffled, more than when it is zeroed or when noise is injected into the MTL. **e**, Decoding accuracy for context identity, plotted with respect to steps since a context switch. Dotted line, chance (0.08, twelve contexts).

### 2.3 A continuous context code in the MTL latent

A key feature of the CB-MTL circuit is that all contexts share a latent space, so novel contexts can be represented in relation to experienced ones. This is what enables a downstream policy to transfer between contexts, and distinguishes this circuit from discrete methods like COIN ^2^. We therefore froze the circuit and measured the latent *z* under 24 novel contexts—rotations and curls within and beyond the training range and drifts at new directions and speeds—and asked whether the latent encoded the relations (such as perturbation family or sign) between these contexts.

Each perturbation family was represented in its own subspace. Rotations were organized along the first principal component while curls were represented along the second principal component (Fig. 3a). Drifts were represented in the subspace spanned by the third and fourth principal components (Fig. 3d).

Location within each subspace was determined by the sign and magnitude of each perturbation; novel contexts interpolated between and extended beyond the training points. Rotations lay along an axis based on angle. Held-out rotations within the training range fell between the training points, and those outside the training range extended the axis, though with compressed spacing (Fig. 3b). Curls were similarly organized along an axis based on the signed gain (Fig. 3c). Drifts formed a series of concentric rings with direction encoded by angle on the ring and speed by radius (Fig. 3e,f).

The latent’s location within this subspace was updated continuously as prediction error accumulated. On trials where the context switched, the state began in the region corresponding to the old context and gradually moved to that of the new one (Fig. 3g,h).

Thus, trained only on next-state prediction, the CB-MTL circuit learns to represent familiar and novel contexts in a shared latent—organized by family and parameter—that is updated as evidence accumulates. We next asked whether a policy receiving this latent learns faster, and whether it can use the structure to compensate for contexts it has never seen.

**Figure 3.**
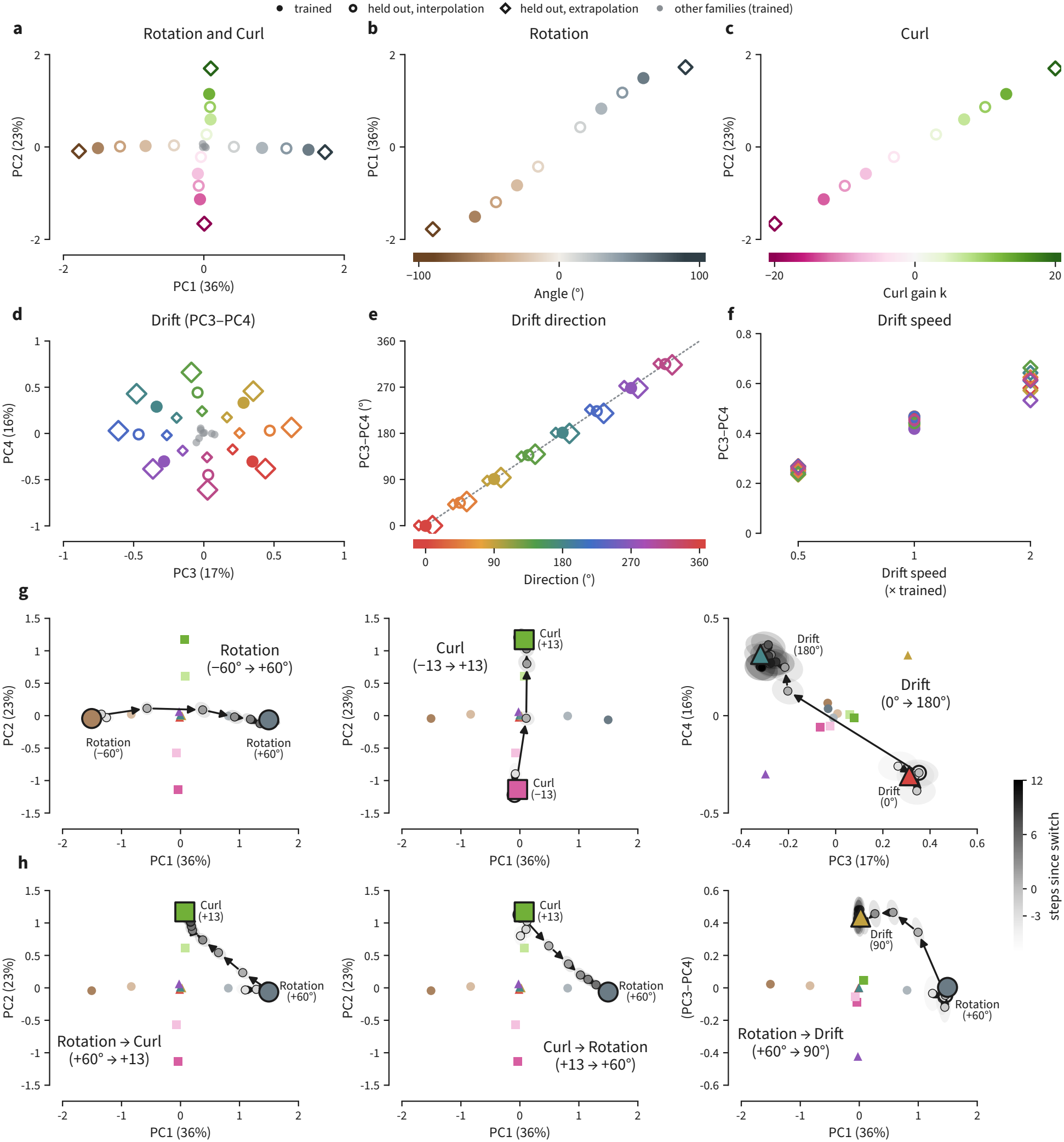
The MTL latent continuously represents perturbations and updates across a context switch. **a–c**, Context centroids in the top principal components of the latent; trained values filled, held-out interpolation open, held-out extrapolation diamond. Rotations and curls each lie on a monotone one-dimensional arrangement ordered by signed magnitude, with held-out values falling in order between their neighbors. **d–f**, Drifts occupy a separate subspace in which direction maps to angle and speed to radius. **g**, Latent trajectory across a within-family context switch, colored by steps since the switch. **h**, Same, across a between-family switch.

### 2.4 Context inference speeds learning through retrieval and transfer

**Figure 4.**
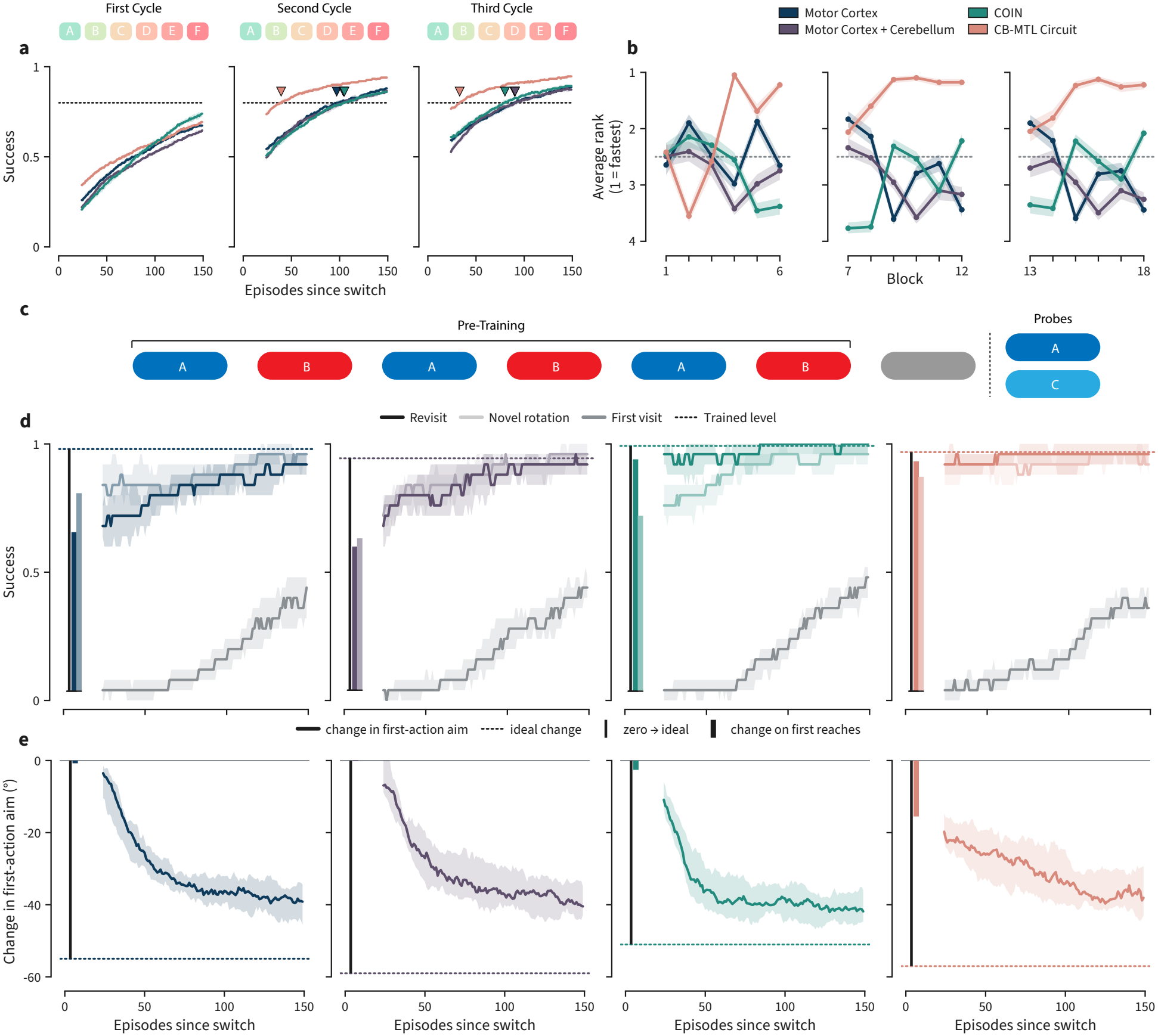
Context inference speeds learning novel contexts and relearning previous contexts. **a**, Success against episodes since switch, one panel per cycle. Each seed’s curve is first averaged over its six contexts; line, median across seeds; band, 95% bootstrap interval of the median. Triangles mark the episode at which each circuit reaches the 80% criterion. **b**, Average rank of the four circuits within each block, 1 = fastest; band, 95% bootstrap interval over seeds. **c**, Retrieval and transfer schedule: alternating A and B blocks, then probes on A (revisit) and C (a novel rotation of the same family). **d**, Success on the revisit and novel probes against first visit, one column per circuit. Line, median across seeds of the smoothed curve; band, inter-quartile range across seeds. **e**, Change in first-action aim over the probe block, same convention; dotted line, the ideal change; bar, the change achieved on the first reaches.

Prior experience can speed learning through three mechanisms, two of which require context. *Competence* is context-independent and improves the performance of a new context regardless of which contexts were originally trained. For instance, learning that running into walls is painful is useful regardless of context. *Retrieval* is context-specific; a policy previously learned for a context can be partially or fully retrieved when the same context or a near similar one is experienced. *Transfer* is graded. It is the estimation of a novel policy based on previous experience. This requires both inferring the current context and relating it to previous contexts already learned. These mechanisms are not mutually exclusive, but architecture limits which ones a circuit can use. MC and MC + CB extract no context, so they can only improve through competence. COIN treats each context as an independent entity and learns no relations between them, so it can use competence and retrieval, but not transfer. CB-MTL represents every context in a shared latent, so it can represent relations among them, thus it can use all three. We therefore predicted that CB-MTL and COIN would benefit from re-exposure to a context, but that only CB-MTL would benefit from exposure to a different context.

We tested this with a cyclic blocked schedule of six contexts, rotations of −60° and 30°, curls of *k* = −13 and *k* = 7, and drifts of 0° and 180°, chosen to span all three families and both signs within each, so that no block trivially solves another. Each context was learned for a block of 150 reaches, switches were uncued, and the six-block cycle was repeated three times, giving three exposures to each context that differ only in how much prior experience of the same and related contexts preceded them. For CB-MTL and COIN, we trained only MC; For MC + CB, we trained CB and MC simultaneously (section 2.1). Comparing learning within a context across exposure reveals which mechanisms operate while comparing across circuits reveals which circuits use which.

No circuit learned quickly on the first block; however, CB-MTL learned quickest from repeated exposures: including on the first exposure to a novel context following prior experiences on other contexts. Median reaches to an 80% success criterion on the first block was 149 for all circuits; 149 indicates the criterion was never reached (Fig. 4a; Extended Data Fig. 3). Nevertheless, averaging across each first exposure (blocks 1-6) the CB-MTL circuit learned fastest (*p* < 10^−5^; Extended Data Fig. 4a). This affect was weak and driven by transfer from previous blocks in the first cycle. Ranking circuits by mean success over the first 50 reaches of each block, CB-MTL led from the 4th block onward: including blocks whose context it had not seen yet (Fig. 4b). The CB-MTL advantage increased as the network was re-exposed to the same context. On the second and third exposures CB-MTL learned fastest and began each block at a higher success rate than any other circuit (Fig. 4a; Extended Data Fig. 4b,c). This suggests that CB-MTL benefited from transfer and retrieval across blocks, but the cycle alone cannot separate transfer from retrieval or competence.

To separate them, we reduced the schedule to two rotations, −60° and 30°, alternated between them for three cycles, followed by a 0° washout, and then probed with either the familiar −60° (revisit) or a novel 60° (Fig. 4c). The three mechanisms predict different trajectories on these probes. Competence alone predicts both trajectories start below the trained level of the −60° and rise. Retrieval predicts the revisit starts near the trained level and the novel context starts low. Transfer predicts both the revisit and novel will start near the trained level.

Both CB-MTL and COIN showed strong retrieval. When re-exposed to the −60° context, both circuits recovered their previous behavior, starting at nearly perfect success despite the two intervening contexts (Fig. 4d). In contrast, MC and MC + CB were both negatively impacted by the intervening contexts. We quantified this as

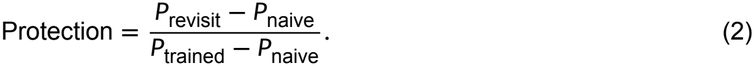

CB-MTL and COIN both began at the trained level, with protection from interference of 1 [0.9-1.02; 25^th^−75^th^ percentile] and 1 [0.9-1] respectively (Extended Data Fig. 4d). MC and MC + CB began below it, at 0.7 [0.5-0.78] and 0.67 [0.44-0.78].

CB-MTL was the only circuit that showed strong performance on the novel 60° context. CB-MTL was the only condition which began the 60° context at near perfect accuracy (Fig. 4d). This suggests that CB-MTL was able to compensate for the novel context based only on prior experience with the −60° and 30° contexts. We quantified this through the first-action aim of the first reaches on the novel block (Fig. 4e)

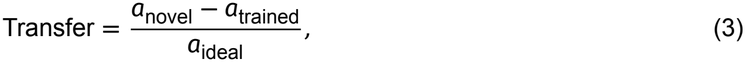

where *a*_novel_, *a*_ideal_, and *a*_trained_ are the first-aims of the novel block, the ideal compensation, and the firstaims of the washout. On those first reaches, MC, MC + CB, and COIN showed no change in aim while CB-MTL compensated by 0.28 [0.19-0.34] of the ideal compensation (Fig. 4e; Extended Data Fig. 4e; *p* < 10^−4^ against each other circuit).

Thus each circuit leverages the mechanisms we predicted its architecture would support. MC and MC + CB improved only through competence while COIN added retrieval. CB-MTL was the only circuit to leverage all three mechanisms. As retrieval is the phenomenon that context inference models of motor learning were developed to explain, we asked whether our circuit can reproduce the three findings that motivated those models—savings after washout, spontaneous recovery, and evoked recovery—and whether MTL was required to do so.

### 2.5 Savings, spontaneous recovery, and evoked recovery require the MTL latent

Three behavioral phenomena motivated context-inference accounts of motor learning. In savings, a perturbation is relearned faster after washout. In spontaneous recovery, adaptation to a prior perturbation re-emerges during error-clamp trials after brief exposure to an opposing one. In evoked recovery, a singletrial of the prior perturbation during the error-clamp recovers its memory. Spontaneous recovery and savings can be reproduced by context-free models ^23,24^. However, evoked recovery, in which a single trial retrieves a memory of a learned behavior, is only explained by context based models ^2^. We asked whether CB-MTL reproduces all three, and whether doing so is dependent on the MTL module. Given the retrieval results above (Fig. 4), we predicted that CB-MTL and COIN would both reproduce all three phenomena, while MC and MC + CB would not.

The savings-washout schedule consists of four blocks: a −60° rotation (P-) for 100 reaches, 60° (P+) for 300, Pagain for 75 as washout, and P+ again for 300 (Fig. 5a). We computed the adaptation index from the first-aim as

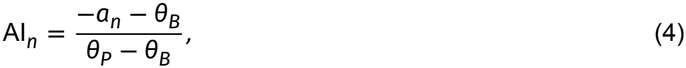

where *a*_*n*_ is the network’s aim direction, measured on the first step, and *θ*_*B*_ and *θ*_*p*_ are the rotations applied during the baseline and perturbation trials: −60° and 60° for the savings schedule. Full compensation for Pgives 0 while the same for P+ gives 1. The adaptation index for all circuits decreased to 0 during the P-block then increased during the P+ block and decreased again during the second P-block. While all circuits adapted during the second P+ block, only CB-MTL and COIN adapted faster on the second exposure than they did on the first. We quantified this through difference between the AI over the first ten reaches of the first and second P+ block (Fig. 5b). CB-MTL and COIN show a benefit of 0.73 [0.58-0.80] and 0.53 [0.40-0.63]. In contrast, MC and MC + CB showed no benefit. Thus the CB-MTL circuit reproduces savings after washout and MTL is necessary to do so.

During the first P+ block of the savings-washout schedule, the adaptation index of MC and MC + CB plateaued at 0.69 ± 0.12 and 0.68 ± 0.09 respectively. This was due to the buffer which draws 15% of samples from earlier blocks. At this point in the task the only preceding context was P-. Because the MC and MC + CB circuits could not learn context-dependent policies. Without a buffer, MC and MC + CB plateaued at 1.00 ± 0.03 and 0.98 ± 0.04.

The CB-MTL circuit reproduced human patterns of spontaneous and evoked recovery. The two paradigms shared a similar schedule. The only difference between the two was that the we inserted a single P+ reach after the tenth channel reach in the evoked recovery schedule. Aside from that, both began with 100 reaches of 0° (P0) followed by P+ for 300 reaches, P-for 50 reaches, then 150 reaches in a channel block. During the channel block, learning was frozen and the agent always moved in a line from the start to the goal— regardless of its actions. Thus preventing either COIN or CB-MTL from correctly updating the context, which stayed near its initial value through out the reach. The initial context for both models was sticky (Methods); with high probability the state (context) of the next trial was the state at the end of the preceding trial. The adaptation index for CB-MTL and COIN increased during the channel block. MC and MC + CB, in contrast, have a steady AI (Fig. 5c). We quantified this through the recovery index

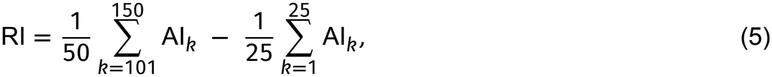

which was 0.82 [0.29-1.34] and 0.97 [0.89-1.07] for CB-MTL and COIN but zero for MC and MC + CB (Fig. 5d). The adaptation index of all four circuits jumped after the evoked trial, but the jump was largest for CB-MTL and COIN 1.47 [1.14-1.62] and 1.70 [1.59-1.77], respectively, compared to 0.14 [0.05-0.29] and 0.28 [0.09-0.47] for MC and MC + CB (Fig. 5f, filled). CB-MTL and COIN were also the only circuits to show a decay over the channel block (Fig. 5f, open).

In both schedules, the policies are frozen, so what changes are the relative weights of the beliefs of the learned contexts. Decomposing the channel block AI by the memory each reach was dealt shows that AI conditioned on each memory remains constant, and that recovery is driven by the reweighting of contexts in the buffer (Extended Data Fig. 5; Extended Data Fig. 6).

The CB-MTL circuit therefore reproduces the three phenomena—savings after washout, spontaneous re-covery, and evoked recovery—that motivated context inference motor learning models, and as predicted MTL was required for reproducing these effects.

**Figure 5.**
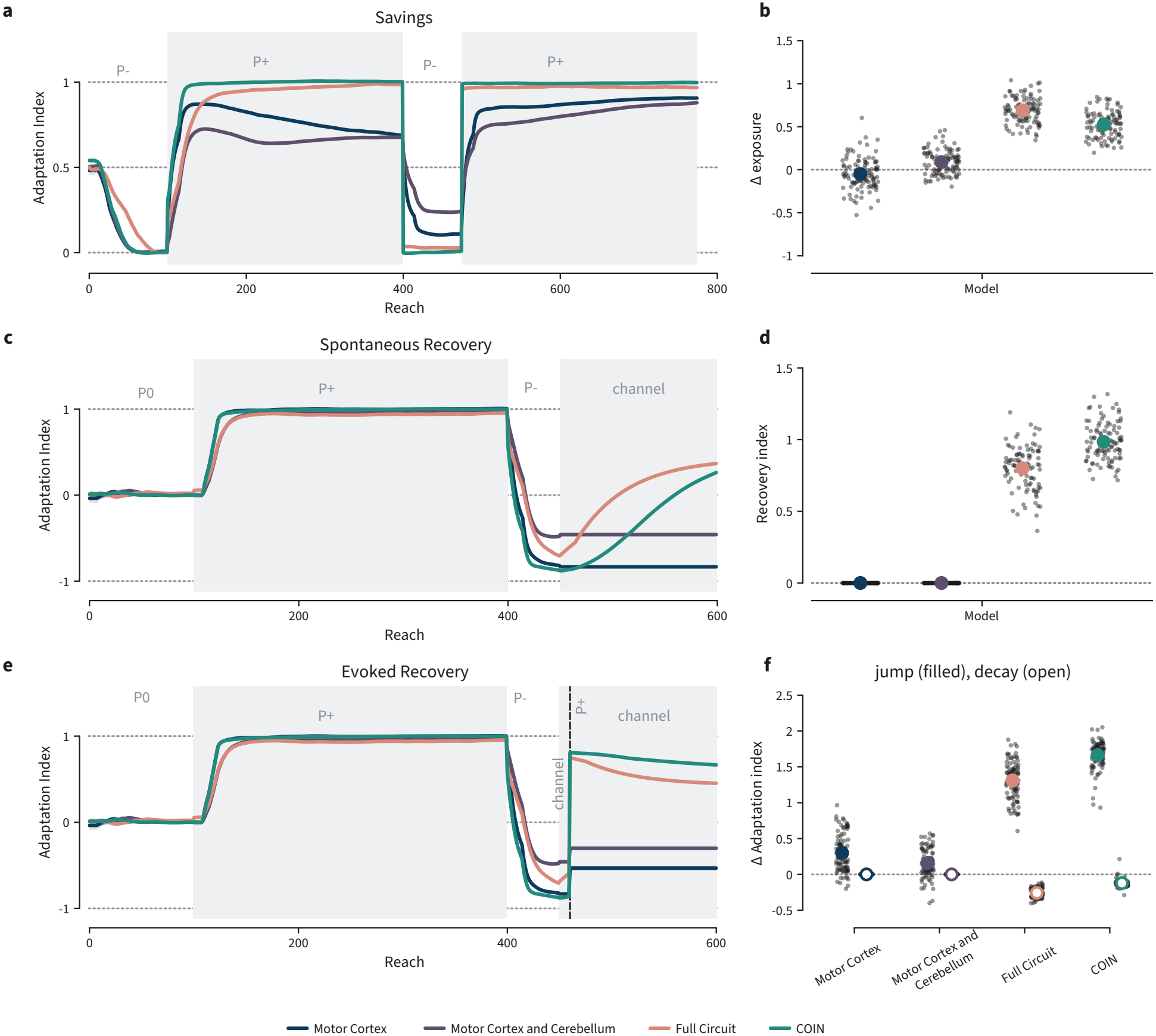
Savings, spontaneous recovery and evoked recovery all require MTL. **a**, Adaptation index (Equation (4)) across the savings–washout schedule. **b**, Re-exposure benefit: mean AI over the first ten trials of the re-exposure block minus the same average on first exposure; dots are networks. **c**, Spontaneous recovery; channel trials withhold prediction error, so no context can be inferred. **d**, Recovery index (Equation (5)). **e**, Evoked recovery; a single P+ trial after ten channel trials. **f**, Filled, the jump in AI immediately after the evoking trial; hollow, its subsequent decay over the channel block

### 2.6 Structured experience enables within-family generalization

Humans can structurally generalize from prior experiences with visuomotor rotations to more quickly learn novel rotations ^1^. The structured representations formed by the network (Fig. 3) and the transfer results above (Fig. 4e) both suggest that the CB-MTL circuit should similarly generalize to novel contexts of perturbation families it previously experienced. Following Braun et al. ^1^, we pre-trained two additional circuits using the curls and drift contexts previously defined. The two circuits differed on whether they were additionally exposed to rotations (±30° and ±90°) or two four random linear transformations. Following next-state prediction training, each policy was trained on four anchor rotations (±30° and ±90°). We then froze all weights of the network and tested generalization to held out contexts of the rotation family between ±90°. For reference, we also compared against COIN to get a baseline of how well a model with retrieval but not transfer would perform.

Linear decoders accurately predicted the angles of novel rotations from the representations of both the rotation-trained and the random linear-trained circuits (Fig. 6a). However, the two differed in their first-angle compensation (Fig. 6b). The rotation-trained circuit compensated nearly perfectly at untrained angles. The slope of its corrections was 0.88 (0.87-0.90; 95% CI). In contrast, the slope of the random-linear trained circuit was half as strong (0.48 [0.46-0.49]). This gap remained even when we allowed both networks to train on novel rotation 60° rotation (Fig. 6c). The rotation-trained circuit had a larger initial return than the random linear-trained circuit and that gap remained throughout training. Thus the CB-MTL circuit produced the same pattern of curriculum-dependent structure learning effects in human motor learning ^1^.

**Figure 6.**
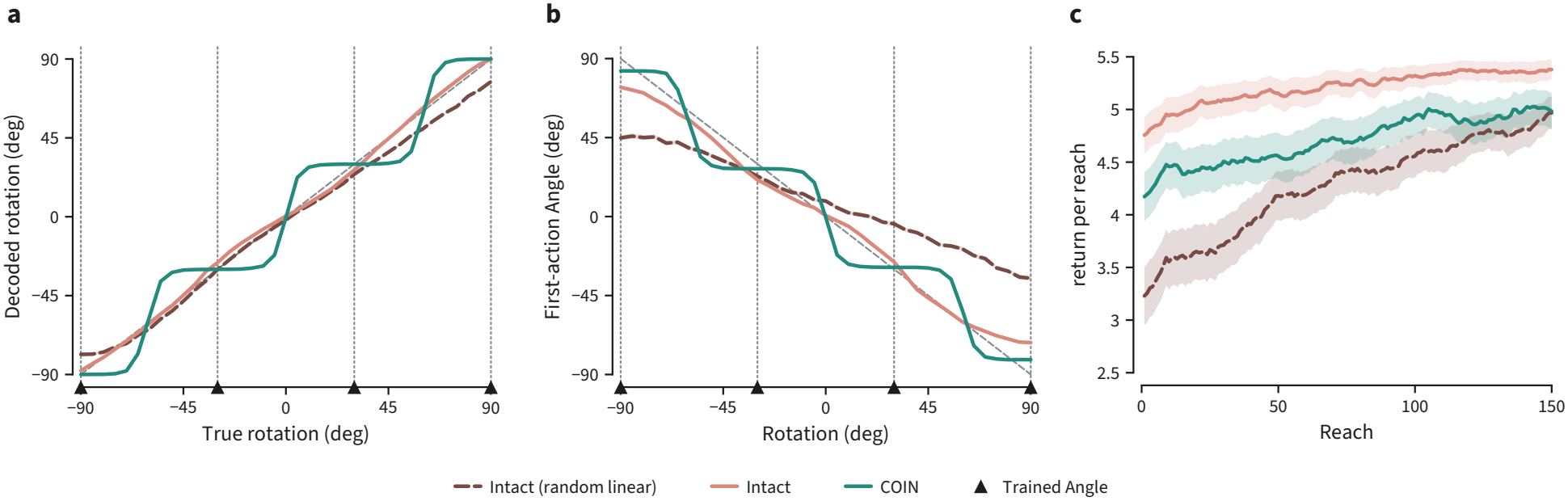
Structured training and MTL enable generalization to novel contexts within a family. **a**, Linear decode of rotation from the latent, trained on ±30° and ±90° and tested across the range. Both circuit variants track the identity line; COIN’s decode is a staircase on the trained values. **b**, First-action aim across the range with no training. The circuit pre-trained on rotations aims near ideal compensation at every angle, while the variant pre-trained on random linear transformations only weakly modulates its aim; COIN is unaffected by the pre-training contrast by construction. **c**, Return per reach while learning a novel rotation.

### 2.7 Inferring where an object is and what it is

The sections above show that the cerebellar prediction error is essential for MTL to infer the latent context. The circuit makes a sharper claim than that, about when the error should matter: it should be needed when an animal’s own actions change what it will sense. The difference between what is sensed and what the cerebellum expects carries the latent information that the MTL cannot directly observe and this information enables MTL to infer the context.

For example, the location of an object with respect to an animal can change either because the object moved or because the animal did, so a change in its bearing or apparent size is not on its own enough to tell which, and only a forward prediction of the animal’s own contribution separates them. The identity of an object does not vary with the animal’s own movement, so there the forward model adds nothing beyond what the animal can already sense. The circuit therefore predicts that the cerebellar prediction error is required to notice that an object has been displaced and dispensable to notice that it has been replaced. Such a dissociation has been reported by Zeidler et al. ^14^. Acute cerebellar stimulation perturbs hippocampal representations and produces a behavioral deficit on an object-location memory task (OLM) while leaving an object-recognition memory task (ORM) intact.

To test whether the CB-MTL circuit could reproduce the dissociation observed in Zeidler et al. ^14^, we place an agent in a circular arena containing two objects (Fig. 7a,b). The agent never has access to its own position, and what it senses of an object degrades with distance and with angle farther away from its heading, so a close reading is exact and a far one is noisy (Methods). The arrangement of the arena is therefore never available at once and must be accumulated across a trajectory. MTL performs that accumulation and passes the inferred latent to cerebellum, whose prediction is conditioned on it at every step.

We follow the two-session design of Zeidler et al. ^14^. The agent explores, after which one object is either moved (OLM) or changed in identity (ORM), and then it explores again. We read the prediction error over the four steps following the change and summarize it as a preference index, PI = *e*_changed_/(*e*_changed_ +*e*_unchanged_), which sits at 0.5 when the change goes unnoticed.

The gap between prediction error for the changed and unchanged object is the analog of the exploratory preference: it measures how much more surprising the altered object is than the unaltered one. In the intact circuit this gap is large and transient in both tasks, widest at the change and closing as the model relearns (Fig. 7c,d) — the circuit is surprised both by a displacement and by a change of identity. Perturbing the cerebellum abolishes the gap for location (Fig. 7e): the two perturbed curves lie almost on top of one another. Critically, this is not a failure to respond to the object that moved. The perturbation raises error on both objects, but it raises error on the unchanged object far more, so the two converge and the contrast is lost. Identity behaves differently (Fig. 7f). The perturbation raises the floor there too, but the unchanged object remains unsurprising and the gap survives nearly intact.

Summarized as a preference index, the dissociation is graded with perturbation strength and is site specific (Fig. 7g,h). For location, noise on the cerebellar output drives the index steadily toward chance, with a fall already measurable at the weakest perturbation tested; corrupting the cerebellar error channel alone produces the same effect, more slowly. Noise delivered directly to the MTL recurrent state leaves the index nearly flat across the full range. Detecting a displacement requires a forward prediction of where the object should appear, which is the cerebellum’s role, and does not require a memory of where it was, which is MTL’s. For identity, the index is spared at every site and every strength tested, remaining near ceiling throughout (Fig. 7h): a shape is not transformed by the animal’s movement, so there is nothing to predict, and a single close reading pins it exactly, so little rests on MTL.

The dissociation does not depend on the particular perturbation strength or the averaging window that we chose (Extended Data Fig. 8). Extending the perturbation well beyond physiological strength and varying the post change window over which the index is computed, the location preference is lost at a lower perturbation strength than the identity preference at every window tested, and the ordering never reverses.

**Figure 7.**
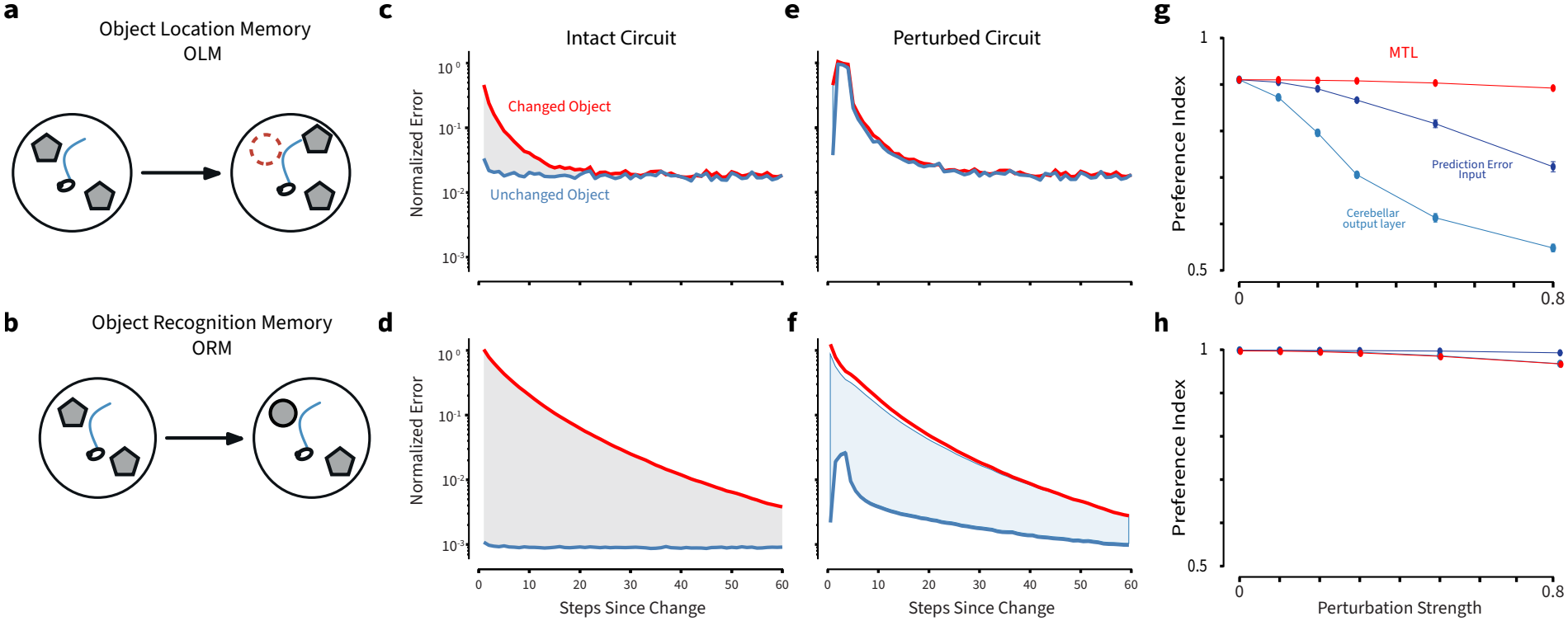
Object location and object recognition memory. **(a, b)** The two paradigms: one object displaced (OLM) or given a new identity (ORM). **(c-f)** Prediction error for the changed and unchanged object following the change, intact and with the cerebellum perturbed; the shaded region is the preference signal. Curves are means over 10 independently trained networks. **(g, h)** Preference index against perturbation strength at three sites, for OLM and ORM respectively; the dotted line marks chance.

The dissociation does not depend on the particular sensory geometry we chose (Extended Data Fig. 9). We retrained the circuit across a sweep over three parameters of the observation model — the angular extent over which bearing is read cleanly, the visual radius, and the whisker reach within which shape is sensed by touching — and repeated the full perturbation protocol at each setting, with three independently trained networks per point. The task remains solvable throughout: the intact preference index stays high for both tasks at every setting (Extended Data Fig. 9a-c), so the costs below are always a fraction of a real preference. Across the range tested, cerebellar perturbation costs object-location memory more than object-recognition memory, and perturbing MTL costs neither (Extended Data Fig. 9d-f). The location cost is essentially flat across the sweep, while the identity cost stays small wherever the whisker reach is shorter than the arena radius — that is, wherever shape must be read from contact rather than being available everywhere at once. The network therefore reproduces the dissociation reported by Zeidler et al. ^14^, and locates its cause in a single asymmetry: a judgement about where something is must be made against a prediction of one’s own movement, and a judgement about what it is need not.

The same asymmetry can be asked of a latent the animal never observes at all: its own position. With a cue in view, the position is reanchored at every step, so an inaccurate forward model can be compensated for before its error accumulates, so cerebellar damage should be invisible. In darkness, there is nothing to reanchor against and the cerebellar prediction becomes the sole estimator of self-motion. The circuit therefore predicts a deficit that is absent in the light and severe in the dark — not because the dark is harder, but because the cerebellum is then the only source of the estimate.

This is what Rochefort et al. ^13^ report. They recorded CA1 place cells in L7-PKCI mice, in which protein kinase C inhibition in Purkinje cells blocked cerebellum-driven plasticity. The animals foraged in a circular arena across five sessions in which a visual cue was present, removed, and restored. In darkness the mutants’ place fields showed reduced stability, coherence and firing rate, and recovered once the cue returned, while wild type fields were largely unaffected.

The simulation reproduces the dissociation in a circuit trained only to predict the next state (Extended Data Fig. 10a). Individual rate maps show that in the intact circuit the map is largely unchanged by cue removal, whereas in the mutant it reorganizes during the unlit sessions and returns once the cue is restored. The three quantities Rochefort et al. ^13^ report behave the same way (Extended Data Fig. 10b): in the light the genotypes are indistinguishable on mean field rate and spatial coherence, and both fall in the dark, but roughly twice as far in the mutant. Between-session stability declines in both as sessions grow further apart, but declines far more steeply in the mutant, and recovers in both once the cue returns. Quantifying the maps directly on four similarity measures gives the same answer (Extended Data Fig. 10c): in the light the two genotypes coincide almost exactly on every measure, and only the dark condition separates them.

Across both experiments the same rule accounts for when the cerebellum is required by the MTL and when it is not. The cerebellum is needed wherever the inference depends on a forward model — where what should be sensed next must be computed from the animal’s own action, such as for an object’s location or for self position in darkness — and dispensable wherever the latent is read directly from the senses or can be continuously re-anchored by an external cue, as for position in the light.

### 2.8 Context inference is prediction error agnostic: reward prediction error enables context inference too

The sections above show that the MTL infers a latent context from cerebellar prediction errors when the context governs how the animal’s own actions change its state: where an object will be sensed, and where the animal itself will be. However, many contexts we experience do not act on position at all. They leave the state unaltered and only change the outcome. In such a setting, the cerebellar prediction error is about value rather than state: a scalar reporting only the difference between the reward received and the reward expected, carrying no information about the state the action produced. The cerebellum carries reward signals as well as sensory ones — reward expectation is encoded in granule cells ^25^, reward prediction errors (RPE) are conveyed by climbing fibres ^26^, and the cerebellum projects directly to reward circuitry ^27^ — so a signal of this kind is available to the MTL by the same route. We therefore ask three things: can the MTL infer the context from a cerebellar RPE alone; is the context it infers represented in an abstract, generalizable form; and does the inference survive removal of the error?

We test this by simulating the serial reversal stimulus–response–outcome (SRO) task of Bernardi et al. ^8^, in which the context does not alter the state but only the outcome an action produces. Stimuli are drawn independently on every trial, so nothing about the next observation can be anticipated, and the only prediction error available to the MTL is a reward prediction error. The task has two uncued latent contexts and four stimuli; each (stimulus, context) pair fixes a correct operant and whether performing it pays, and at the switch, the correct operant reverses for all four stimuli while the reinforcement contingency reverses for two of them, so that operant and reward remain orthogonal (Fig. 8a).

The geometry of the MTL hidden state shows that context is both decodable and abstract. Averaging activity within each of the eight (context, stimulus) conditions and projecting those means onto their first three principal components, separates the two contexts along the leading component while preserving the arrangement of the four stimuli within each (Fig. 8b). Distances measured in the full space, not in the projection, give the same picture: conditions sharing an operant lie closest whereas conditions differing in operant lie further apart, and the two contexts are separated by more than either. The same stimuli in the two contexts are joined by lines that run parallel, which is the geometric form of the parallelism score. Fitting a linear context boundary to six of the eight conditions, leaving out both conditions of one stimulus, classifies that held-out pair correctly, showing the abstract nature of inference.

Context becomes linearly decodable from the MTL hidden state, and after a switch it is recovered within a few trials (Fig. 8c). It depends on the reward prediction error exactly as the context depended on the state prediction error above: replacing *δ*_*t*_ with zero at test or shuffling it across sequences leaves decoding at chance at every lag from the switch (Fig. 8c). The augmented control arm, which receives the previous reward directly, has a second route to the outcome and is unaffected by the same ablations, though it recovers the context to a lower level than the intact main arm (Extended Data Fig. 11). The reward-prediction error is therefore not merely correlated with the context code; it is the only evidence MTL has for it.

The representation of the circuit here matches the recordings in the hippocampus. Applying the measures of Bernardi et al. ^8^ to the MTL hidden state, all three task variables are decodable, but only context is abstract (Fig. 8d). Context reaches near-ceiling cross-condition generalization while previous value and previous action fall far below it, and the same separation appears in the parallelism score, where context is near one, action is negative and value sits at zero (Extended Data Fig. 11). A context decoder trained on some stimuli therefore reads context correctly on stimuli it has never seen, while a value decoder does not: a switch reverses which responses pay for only half the stimuli, so a value decoder trained in one context transfers poorly to the other — hence a low cross-condition generalization alongside a parallelism score of zero. This abstraction is not bought at the cost of dimensionality. The shattering dimensionality of the MTL hidden state is near one, meaning a linear decoder recovers nearly all 35 balanced dichotomies of the eight conditions (Extended Data Fig. 11). The model thus matches the hippocampal recordings on both counts Bernardi et al. ^8^ report — an abstract context code held within a high dimensional geometry.

The same pattern holds throughout the circuit, and only while the reward-prediction error is intact (Extended Data Fig. 11). In the main arm, context, previous value and previous action are each recovered at nearceiling accuracy from the MTL hidden state and from both cerebellar layers. The augmented control reaches comparable accuracy in the cerebellar layers but falls well below it in the MTL hidden state, where context in particular is only weakly decodable. Replacing *δ* with zero or shuffling it drops context and previous action to chance in every layer, while previous value remains partly decodable, since under this design reward is fixed by stimulus identity alone for half the stimuli and requires no inference about context (Extended Data Fig. 11a). The abstraction measures separate the variables sharply. In the main arm, context reaches high cross-condition generalization and a parallelism score close to one, while previous value and previous action fall near zero or below on both (Extended Data Fig. 11b,c). The abstract format is strongest in the MTL hidden state and weakens through the cerebellar layers, with the context parallelism score falling from near one in MTL to intermediate values in layer 2 — consistent with the cerebellum inheriting the context code rather than constructing it. The augmented control represents context weakly, with decoding below the main arm, and its abstraction both lower and far more variable across networks. Shattering dimensionality behaves differently from the other measures: when the prediction error is perturbed it falls only partway and remains well above chance (Extended Data Fig. 11d). The ablated network therefore still carries a high dimensional representation and can still separate arbitrary dichotomies; what it loses is the particular variables and the abstract axis along which they were organized.

**Figure 8.**
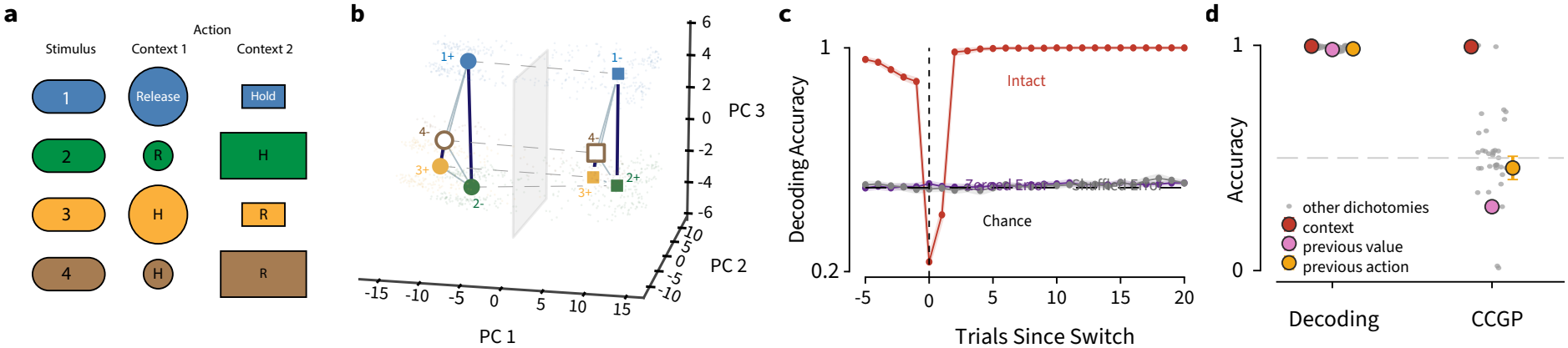
Context inference from a reward prediction error. **(a)** The serial-reversal SRO task: four stimuli, two uncued contexts, each pair fixing a correct operant and whether it pays. **(b)** The MTL hidden state separates the two contexts while preserving the same stimulus arrangement within each. Circles, context 1; squares, context 2; color gives stimulus identity and the sign whether the correct operant pays. Dark lines join conditions sharing an operant, pale lines conditions differing in it, and grey dashed lines the same stimulus across contexts. The plane is a linear context boundary fitted to six conditions in these coordinates, with both conditions of stimulus 4 held out (hollow); it classifies that held-out pair correctly. **(c)** Context decoding accuracy relative to a switch, for the main arm, with the prediction error intact, zeroed and shuffled, averaged across 10 seeds. **(d)** Decoding and cross-condition generalization for each task variable, against all 35 dichotomies.

## 3 Discussion

Context inference and structure learning are crucial to our ability to learn context-dependent behaviors across motor and non-motor tasks. Yet, little is known about how these computations are performed in the brain or whether a shared circuit might support their use in motor and cognitive tasks. We asked whether a shared cerebellar-medial temporal lobe circuit could support context inference and structure learning. We found that training a circuit consisting of a cerebellar forward model and a recurrent medial temporal lobe module to predict its next-state was sufficient for enabling context inference and structure learning. The circuit learned to represent contexts—trained and novel—in a continuous low-dimensional latent space, where distinct subspaces represented the different types of contexts and location within a subspace was determined by the parameter of the perturbation. This structured context representation enabled a motor cortex model receiving this input to generalize across contexts, and reproduce context-dependent learning phenomena. The same circuit was able to extract context in a non-motor task and its medial temporal lobe module reproduced patterns of hippocampal activity observed in the same task. Further, we show that when the same circuit is applied to three memory tasks, it reproduced the pattern of cerebellar-dependence of each: including the sensitivity of hippocampal place cell activity to cerebellar stimulation in the dark. Thus we propose the cerebellar-medial temporal lobe as a general circuit for context inference and structure learning in the brain.

### 3.1 Relation to other models

Our model is primarily motivated by COIN ^2^ and seeks to reproduce the same context-dependent features that motivated it: savings, spontaneous recovery, evoked recovery, and context-dependent retrieval. However, while COIN provides a normative account of these phenomena, it does not provide a mechanistic account of how such computations can be performed in the brain. Our circuit does; it reproduces the core phenomena of COIN while also providing a model for how such computations can be performed by neural circuits in the brain. An additional advantage of our circuit model is that it can generalize to novel contexts through structure learning. This allows our model to reproduce structure learning-dependent features of human motor learning such as experience dependent effects in learning visuomotor rotations ^1^.

Previous models have shown that hallmark features of entorhinal activity emerge when a recurrent network is trained on path-integration. In path-integration models, the network receives the animal’s velocity as an input and is trained to predict the next position of the animal. The network must therefore learn to integrate the provided velocity signal. In doing so, these models produce grid cell codes ^28,29^. The Tolman-Eichenbaum Machine (TEM) extends this to non-spatial structure and shows how structure learning can emerge from such a model ^30^. Its entorhinal module integrates actions to build a structural code of the environment and the hippocampal module binds this code to sensory observations. Because the structural code is separated from the sensory code, the network can generalize to environments that share similar structure but disparate sensory mappings. Our circuit differs from these prior models in what is integrated. In path integration and in TEM, the integrated input is the animal’s velocity or action, both veridical. In our model, the integrated input is the prediction error of a learned forward model. The prediction error accounts for the features left unexplained by the forward model and it can be integrated to infer the hidden latent cause of the model’s own discrepancy. In this sense, the prediction error can be thought of as an abstract velocity in the space of latent causes rather than a physical velocity through the world. Upon this change, the circuit reproduces cerebellar-dependent features of MTL activity (Fig. 7; Extended Data Fig. 10), the abstract geometry of hippocampal activity (Fig. 8), and context-dependent learning and structure learning in motor control (Fig. 2; Fig. 5; Fig. 6).

Previous work has also shown that features of hippocampal activity can be reproduced by training a recurrent network on next-state prediction. Jensen et al. ^31^ trained a recurrent network to plan, under a predictive auxiliary loss, and found that the network produced hippocampal-like replay during planning. Similarly, Chen et al. ^32^ showed that training a recurrent model of the medial temporal lobe to predict next-state produced place cells in their CA1 module. Our model belongs to a similar prediction family; however, it differs in where the prediction is performed. In both Jensen et al. ^31^ and Chen et al. ^32^ the prediction is performed by the recurrent network itself (i.e. medial temporal lobe) while in our model the medial temporal lobe provided a latent signal that aided next-state prediction in the cerebellum. This difference was important for explaining patterns of cerebellar dependence in our memory tasks.

### 3.2 Role of the medial temporal lobe in motor control

McDougle et al. ^33^ tested a patient, L.S.J, with severe bilateral MTL damage on four motor learning tasks. While L.S.J adapted to perturbations, she was slower to de-adapt following removal of the perturbation and demonstrated weaker savings than age-matched controls ^33^. In particular, L.S.J showed no learning benefit from re-exposure to the same perturbation. These results suggest impaired context-dependent retrieval and updating following MTL ablation. Our model produces the same pattern. In our model, savings has three sources: competence, retrieval, and transfer. Competence is unaffected by MTL ablation while retrieval and transfer are. Without MTL the model never acquires context-dependent policies, so the policy learned under the perturbation gets overwritten by intervening contexts and is unavailable on re-exposure. The deficit therefore has a specific source: MTL infers context from the cerebellar prediction error, and without that inference there is nothing under which a policy can be stored separately and recovered.

### 3.3 Role of cerebellar-medial temporal lobe interactions

While typically studied in the context of motor control, the cerebellum’s outputs reach well beyond motor cortex. Transneuronal tracing reveals reciprocal connections between the cerebellum and prefrontal cortex ^34^, parietal cortex ^35^, and medial temporal lobe ^11^. Its activity reflects non-motor variables, including expected reward ^25,36^, reward prediction error ^26^, and accumulated sensory evidence ^37^, and cerebellar damage produces cognitive and affective deficits ^38^. Despite this evidence of interaction, its function remains unclear, as does which features of neural activity and behavior depend on it. Object-location memory, for instance, is cerebellar-dependent while object-recognition memory is not ^14^. Our model suggests that both findings follow from a shared cerebellar computation, forward model prediction ^17–19^, coupled with the known anatomy ^11^. Given these two constraints, the model produces the object-location and object-recognition dissociation (Fig. 7), the cerebellar dependence of place cells in the dark (Extended Data Fig. 10), and the abstract geometry of hippocampal activity (Fig. 8). In each case, our model predicts that medial temporal lobe activity and behavior depend on the cerebellum when the consequence of an action must be inferred from prediction error, but not when it is directly provided by the input.

### 3.4 Limitations and future directions

The primary limitation of this work is that the circuit reproduces previously reported data but does not itself test new predictions. Every behavioral and neural result presented here is a reproduction of a published finding. Our model instead offers a shared mechanism for explaining these previous results which enables testable predictions about the cerebellar–medial temporal lobe circuit. For example, our model predicts that context inference during the SRO task is cerebellar-dependent, and that cerebellar disruption during the serial-reversal task will impair context inference. It also predicts that structural generalization to a family of visuomotor rotations or velocity-dependent curls is medial temporal lobe-dependent, and that the medial temporal lobe represents the structure of these perturbations in a continuous latent space.

Another limitation of this study is that the circuit is schematic rather than an anatomically and physiologically detailed model. Our circuit collapses entorhinal cortex and hippocampus into a single recurrent module and ignores cerebellar architecture, such as the fan-in fan-out connectivity. We also made many simplifying assumptions during network training, for example that the weights of cerebellum and medial temporal lobe are frozen while motor cortex learns the navigation task. None of these simplifications bears on the main conclusions of this work—which tests whether these components are sufficient for context inference, but each limits what conclusions can be derived from the results presented here. For example, the current work cannot answer how hippocampus and entorhinal cortex differentially contribute to context inference, how the sparsity of Purkinje cell activity affects it, how short-term or behavioral-timescale plasticity could further support it, or how context inference is affected by the loss of specific cerebellar, entorhinal, or hippocampal cell types. Answering each requires extending one component of the present circuit.

A future direction is to use the inferred context to aid memory retrieval. Recent circuit models propose that the entorhinal–hippocampal system maintains a fixed set of memory addresses built from grid cell activity, so that storing a memory means assigning it to an address and retrieving one means letting a cue settle onto the nearest ^39^; in key–value terms, the hippocampus holds the addresses and the neocortex the content ^40^. Keeping the addresses fixed allows recall to degrade gradually rather than suddenly collapse ^41^, but it guides retrieval by superficial sensory resemblance. However, two situations that look alike might demand opposite responses, and the experience that helps may share little perceptually, causing the sensory similarity to settle on the wrong address. For example, recall in post-traumatic stress disorder has this character: the sound of fireworks resembles gunfire enough to recall a traumatic memory, even though nothing else about the situation resembles the one that formed it. Our circuit provides an alternative route: indexing retrieval by the inferred latent context rather than by sensory similarity predicts spontaneous, reversible remapping in an environment where no sensory cue changes ^42^. This also generates a prediction about the dynamics of information flow between circuits, which should be visible in neural recordings: a change in the cerebellar prediction error should precede the switch in the hippocampal map rather than follow it.

### 3.5 Conclusion

We have shown that a circuit built from a cerebellar forward model and a recurrent medial temporal lobe, trained to predict next state, infers latent context. The medial temporal lobe represents contexts, both trained and novel, in a continuous latent space: organizing contexts by their structural relations. A motor cortex receiving this latent as an input reproduces context-dependent features of motor learning, such as savings, evoked and spontaneous recovery, and compensation for perturbations it never encountered. The same circuit reproduces the cerebellar dependence of hippocampal activity and behavior across three memory tasks. Cerebellar–medial temporal lobe interaction is therefore a candidate locus for context inference and structure learning in the brain.

## 4 Methods

### 4.1 Environment Dynamics

The agent is a point mass moving in a continuous two-dimensional square arena, [−1, 1]^2^. The agent navigates this environment by producing two-dimensional forces, *F*_*t*_ ∈ [−1, 1]^2^. On each step, the agent’s force is corrupted by signal-dependent noise

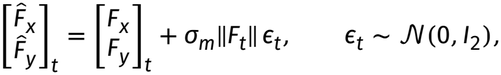

where σ_*m*_ = 0.28 is the motor-noise scale and 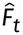 is the corrupted action. The corrupted action is transformed by the current context *k* and integrated as

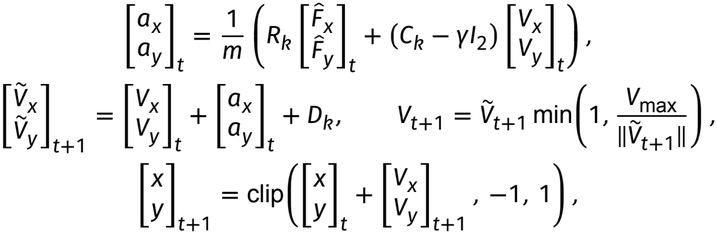

where *m* = 32, *γ* = 1 and *V*_max_ = 0.1 are the agent’s mass, the damping coefficient and the maximum speed, respectively. Collisions with the walls are inelastic: if a step would carry the agent past a wall, its position is clipped to the wall and the velocity component normal to that wall is set to zero. The remaining parameters, *R*_*k*_, *C*_*k*_ and *D*_*k*_, are context-dependent and govern rotation, curl and drift, respectively:

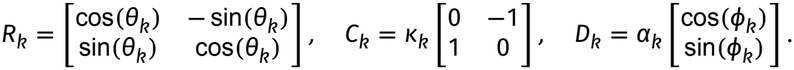

Each context is defined by a unique combination of (*θ, κ, α, ϕ*). In the default pre-training task (detailed below) we train the model on 12 contexts spanning the three perturbation families: rotations of (±30°, 0, 0, 0) and (±60°, 0, 0, 0); curls of (0, ±7, 0, 0) and (0, ±13, 0, 0); and drifts of (0, 0, 0.16, 0°), (0, 0, 0.16, 90°), (0, 0, 0.16, 180°), and (0, 0, 0.16, 270°). These values were chosen to match effect size, 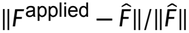, across perturbation families.

### 4.2 CB-MTL Architecture

The CB-MTL circuit consists of two networks, a cerebellar forward model (CB) and a recurrent network standing for the entorhinal cortex and medial temporal lobe (MTL). Both operate on the agent’s observation, *o*_*t*_ = [*x, y, V*_*x*_/*V*_max_, *V*_*y*_/*V*_max_]^⊤^ ∈ ℝ^4^. On each step, CB predicts the next observation from the current observation, the current action and the context latent *z*_*t*_. MTL computes *z*_*t*_ from CB’s prediction error, the previous action, and the previous observation. Both CB and MTL receive the clean intended action *F*_*t*_ not the noisy action 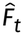.

#### Cerebellum

Cb is a feedforward network with two hidden layers of 256 rectified linear units (ReLU),

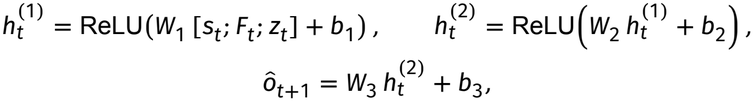

where [⋅ ; ⋅] denotes concatenation. Its input is the 4-dimensional state, the 2-dimensional action and the 32-dimensional latent, 38 dimensions in all. CB predicts the next state (ô*t*) and its error can be quantified using the prediction error,

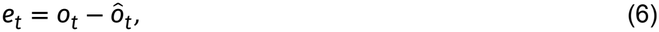

which compares the observed state with the prediction CB made on the previous step.

#### Medial temporal lobe

MTL is a Gated Recurrent Unit (GRU) with 256 hidden units. Its input on step *t* is the previous observation, the previous action, and the current prediction error: *x*_*t*_ = [*o*_*t*−1_; *F*_*t*−1_; *e*_*t*_] ∈ ℝ^10^. Its hidden state is updated as:

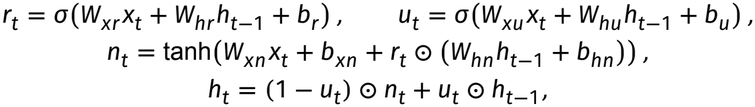

where σ is the logistic function and ⊙ is element-wise multiplication. The latent is a noisy readout of the hidden state,

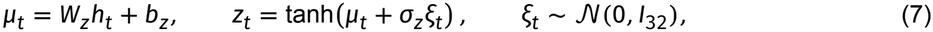

where σ_*z*_ = 0.3 is the noise scale. Noise is applied only during training as a regularization technique.

**Table 1.**
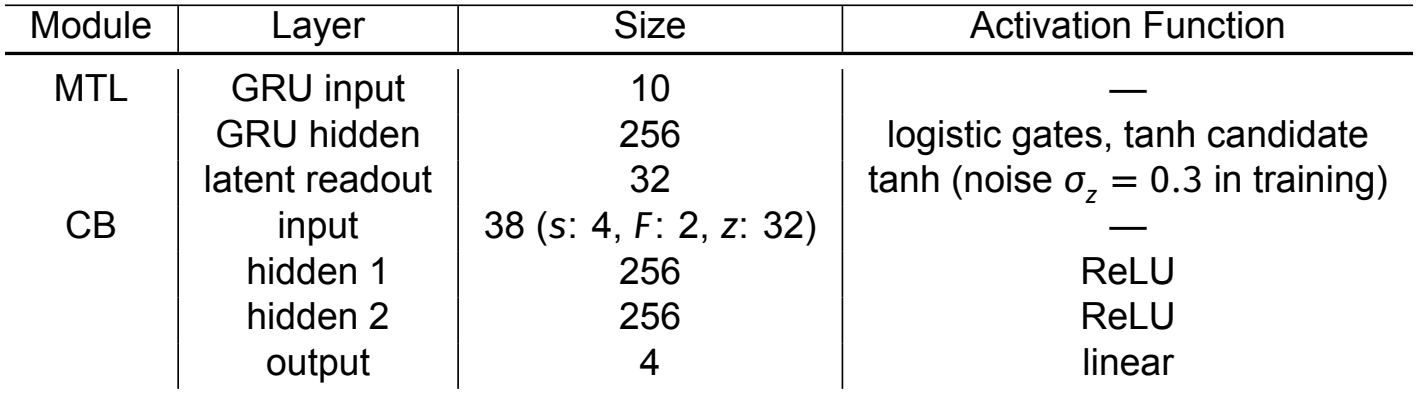
Circuit Specs.

| Module | Layer | Size | Activation Function |
|---|---|---|---|
| MTL | GRU input | 10 | — |
|  | GRU hidden | 256 | logistic gates, tanh candidate |
|  | latent readout | 32 | tanh (noise *σ*<sub>*z*</sub> = 0.3 in training) |
| CB | input | 38 (*s*: 4, *F*: 2, *z*: 32) | — |
|  | hidden 1 | 256 | ReLU |
|  | hidden 2 | 256 | ReLU |
|  | output | 4 | linear |

### 4.3 Next-state prediction training

The circuit was trained on sequences of *T* = 128 steps generated by the dynamics detailed above. Each sequence began with zero velocity at a position drawn uniformly from [−0.5, 0.5]^2^ in a context drawn uniformly from the training set. On every subsequent step, the context switched with probability 0.1 to a different context drawn uniformly from the remaining contexts in the training set. Context blocks therefore had geometrically distributed lengths, with a mean of 10 steps before truncation by the end of the sequence.

Because the optimal actions of a goal-reaching policy are context-dependent, training the model to perform next-state prediction on actions from a trained policy can leak information. To avoid this, we instead sampled sequences of actions from a randomly parameterized Gaussian Process. For each sequence we drew a Matérn smoothness 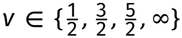 a length scale *l* ∼ LogUniform(0.3, 3) steps, a saturation gain *g* ∼ *U* (0, 5) and an amplitude scale *c* ∼ *U* (0.6, 1.8). Each action component was then generated independently as

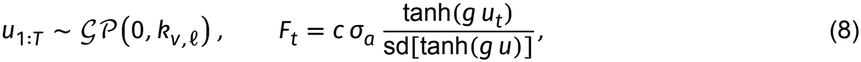

where *k*_*v,l*_ is the Matérn covariance function with unit variance, sd[tanh(*gu*)] is the standard deviation of tanh(*gu*) for *u* ∼ *N* (0, 1), and 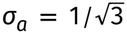. As *g* → 0, the actions are Gaussian with standard deviation *c* σ_*a*_; at large *g*, they are close to binary. The base scale σ_*a*_ matches the standard deviation of a uniform variable on [−1, 1], which approximates the output range of the controller’s squashed policy.

The default circuit was trained on all 12 contexts. For the zero-shot experiment, we trained two further circuits whose rotation family was replaced while the curl and drift families were kept. In the *rotation*-trained circuit, the four rotations were *θ*_*k*_ ∈ {±30°, ±90°}. In the *random linear* -trained circuit, the four rotations were replaced by four random linear transforms of the action, *A*_*k*_ = *I*_2_ + *E*_*k*_, where *E*_*k*_ was drawn uniformly from the sphere 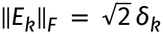. The radii were set so that each transform matched the effect size *δ*_*k*_ of one of the structured rotations. The random transforms were sampled once, from a fixed seed, and shared across circuit seeds.

#### Circuit Training

MTL and CB were trained jointly to predict the next state. The loss was the mean squared prediction error, with velocity weighted more heavily than position, plus a rate cost on the latent channel:

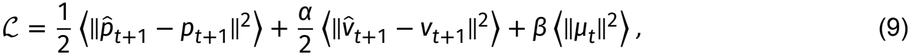

where *p* and *v* are the position and scaled velocity components of *o*, ⟨⋅⟩ denotes the average over sequences and time steps, *α* = 3 and *β* = 10^−3^. We increased the relative weight of the velocity prediction error to account for the next position is highly correlated with the previous position thus the network can achieve a low position error simply by copying its inputs.

Each circuit was trained for 256 epochs on a fixed set of 8,192 sequences, in mini-batches of 256 sequences. Gradients were computed by backpropagation through time over the full sequence. We used AdamW with a learning rate of 10^−3^ and weight decay of 10^−4^. The learning rate followed a cosine schedule with warm restarts, with a first cycle of 50 epochs, a cycle length that doubled after each restart, and a minimum learning rate of 10^−5^. Weights were initialized with PyTorch defaults. Prediction error was monitored on 300 held-out sequences.

### 4.4 Reaching Task

Each episode consists of a single reach. The start and goal were drawn independently and uniformly from the arena, and pairs closer than 0.5 were redrawn. The agent started at rest. Each reach ended when the agent came within 0.1 of the goal, or after 30 steps. The agent received a dense reward

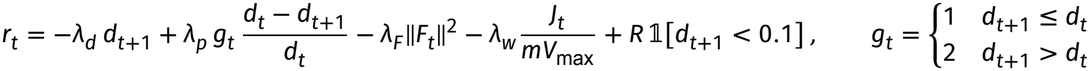

where *d* is the distance to the goal, 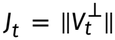 is the impulse of any wall contact on that step (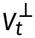 being the velocity removed by the wall), *λ*_*d*_ = 0.3, *λ*_*p*_ = 1, *λ*_*F*_ = 0.1, *λ*_*w*_ = 1/32, and *R* = 5 is the success bonus. The first term of this reward encourages closing the absolute distance to the goal. The second term encourages progress towards the goal. The third term discourages large actions. The fourth term punishes wall collisions. Finally, the fifth term is a large success bonus whenever the agent is sufficiently close to the goal.

#### Action Selection

The action selector (MC) is a soft actor-critic agent ^43^. Its observation is the agent’s position, velocity and goal, [*x, y, V*_*x*_, *V*_*y*_, *x*_*g*_, *y*_*g*_]. Based on the circuit configuration (MC + CB and CB-MTL) the controllers inputs is appended with two additional inputs. The first is MTL’s latent, *z*_*t*_ (32 dimensions).

The second is CB’s second hidden layer, 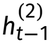, mapped to 32 dimensions by a fixed random projection *p* ∈ ℝ ^256×32^. *p* has unit-norm columns and is drawn once per run.

The actor and both critics are feedforward neural networks with two hidden layers of 256 ReLU units. The actor outputs the mean and log standard deviation of a Gaussian. The log standard deviation is passed through tanh and rescaled to [−5, 2]. The action is a sample from this Gaussian, squashed by tanh into [−1, 1]^2^. Each critic takes the observation and the action and outputs a scalar *Q*-value. Target critics were updated by Polyak averaging such that the target parameters were given by

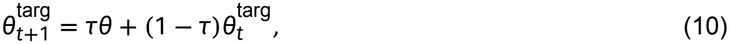

where *τ* = 0.005 is the smoothing parameter: chosen to be small to minimize the variance of this update. We used a discount factor of *γ* = 0.99. The entropy temperature was tuned automatically ^43^ towards a target entropy of −2, starting from a temperature of 1. Actor, critics and temperature were optimized with Adam at a learning rate of 3 × 10^−4^. Gradients of the actor and the critics were clipped to a norm of 1, and all networks used PyTorch’s default initialization. One gradient update was made per environment step, with mini-batches of 256 transitions. Until the replay buffer held 256 transitions, actions were drawn uniformly from [−1, 1]^2^. In configurations with the circuit, MTL and CB were frozen.

#### Replay Buffer

Transitions were stored in a replay buffer indexed by episode, which held the 500 most recent episodes. Sampling weighted each episode by its age *a*, in episodes, as exp(−*a*/40), mixed with a 2% uniform floor over episodes. Each episode’s weight was split equally over its transitions, so an episode’s total sampling weight does not depend on its length. In addition, each mini-batch was split between the block currently being trained and all earlier blocks. The current block received 85% of the sampling weight and earlier blocks 15%, with the recency weighting applied within each part.

#### MTL initial state

MTL’s hidden state was carried from one reach to the next. Each reach began with the hidden state at the end of the previous reach. Each reach’s final hidden state was also stored in a bank. With probability 0.02 per reach, the carried state was replaced by a state drawn uniformly from this bank, so that the start state drifts towards contexts in proportion to how often they have been visited. This plays the role of the sticky transition prior in COIN ^2^. At the start of every reach, MTL’s other inputs (the previous state, the previous action and the prediction error) were reset to zero, as at the start of a training sequence.

We compared four circuits. All circuits share the same reward, buffer, training schedule, and controller. They differed only in which inputs MC received.

#### Motor Cortex (MC)

This circuits contains only MC. Thus MC observed only [*x, y, V*_*x*_, *V*_*y*_, *x*_*g*_, *y*_*g*_]. It has no access to a forward model nor to context inference.

#### Motor Cortex + Cerebellum (MC + CB)

MC with a forward model, through CB, but no MTL. CB has the architecture described above, but its latent input is fixed at zero, so it is a single context-blind forward model, Ŝ _+1_ = CB(*s*_*t*_, *F*_*t*_, 0). Unlike in the CB-MTL circuit, CB was not pretrained. It was initialized at random and trained alongside MC, to predict the next state from replayed transitions. It used the circuit-training loss without the rate term, Adam at a learning rate of 10^−4^, gradients clipped to a norm of 1, and one update per environment step on the same mini-batch as MC’s update. MC observes CB’s second hidden layer through the fixed random projection *p*, as in the CB-MTL circuit. This model therefore tests whether a cerebellar forward model, without a context representation, is sufficient for structure learning.

#### Cerebellum-Medial temporal lobe (CB-MTL)

MC with the pretrained, frozen MTL and CB, observing both *z*_*t*_ and the projected CB hidden state.

#### COIN

MC with an ideal Bayesian observer in place of the circuit, as a finite-hypothesis version of the context inference model of Heald et al. ^2^. The observer’s hypotheses are the contexts the agent is trained on in that schedule. It is given each hypothesis’s true dynamics, which gives an exact likelihood for each context. Steps on which this relation does not hold were treated as carrying no evidence: a wall contact, a step that reached the speed cap, or a zero action. Between steps, the posterior evolves under a sticky transition prior,

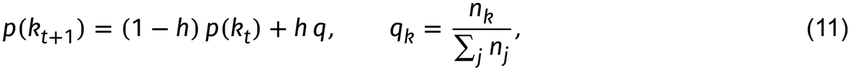

with a hazard of *h* = 10^−3^ per step. Here *n*_*k*_ accumulates, at the end of every reach, that reach’s final posterior probability of context *k*, so the observer jumps towards contexts in proportion to how often they have been visited. Before any reach has been completed, *q* is uniform over the other contexts. The posterior is carried across reaches and blocks, and the observer is never told when the context switches. MC observes the posterior-weighted mixture of fixed context codes, *c*_*t*_ = ∑_*k*_ *p*_*t*_(*k*) *e*_*k*_, where the *e*_*k*_ are unit-norm random vectors in ℝ ^32^, drawn once per run. The null field’s code is zero. The code therefore has the same width as *z*_*t*_ and, at certainty, is a fixed point per context.

### 4.5 Training Schedules

Contexts were presented in blocks of consecutive reaches. The schedules below differ based on which contexts were used, how long the blocks were, and what followed training.

#### Cycle

The default schedule cycled through six contexts, two from each family: rotations of −60° and 30°, curl fields with *κ* = −13 and 7, and drifts in the directions 0° and 180°. Each block was 150 reaches long, and the cycle was repeated three times, giving 18 blocks and 2,700 reaches. Savings were measured by comparing learning in a context’s later blocks with learning in its first.

#### Savings probe

To separate context-specific retrieval from transfer within a family, some cycles were followed by a washout and a set of probe blocks. The washout was a 150-reach block in a neutral, unperturbed, context (*R*_*k*_ = *I, C*_*k*_ = *D*_*k*_ = 0), which removes any carry-over from the last training block. Three probe blocks of 150 reaches followed. Each probe was run as a separate branch from the same end-of-washout state, so both probes started from identical experience and networks. The first probe was a revisit of the −60° context, and the second probe was a novel 60° context.

#### Motor learning paradigms

We reproduced three paradigms from Heald et al. ^2^, using rotations of ±60° as the perturbations P^+^ (60°) and P^−^ (−60°). For these schedules, the agent’s context set was the four table rotations, and COIN’s hypotheses also included the neutral context (P^0^). The block sequences are given in Table 2.

**Table 2.** Block sequences of the COIN paradigms. P^0^: no perturbation; P^+^: 60° rotation; P^−^: −60° rotation.

| Paradigm | Blocks (reaches) |
|---|---|
| Spontaneous recovery | P<sup>0</sup> (100), P<sup>+</sup> (300), P<sup>−</sup> (50), channel (150) |
| Evoked recovery | P<sup>0</sup> (100), P<sup>+</sup> (300), P<sup>−</sup> (50), channel (10), P<sup>+</sup> (1), channel (140) |
| Savings | P<sup>−</sup> (100), P<sup>+</sup> (300), P<sup>−</sup> (75), *P*<sup>+</sup> (300) |

In the savings paradigm, the baseline and washout blocks were the trained −60° rotation rather than the neutral context, because the circuit was never trained on the neutral context.

#### Channel reaches

In a channel block, the environment dynamics were replaced by a clamp that moved the agent straight towards the goal at the maximum speed, regardless of the action. During the channel block, the weights and buffer were both frozen. Because the dynamics changed, neither CB-MTL nor COIN could correctly update their beliefs.

Throughout each paradigm, the expressed adaptation was also read out after every reach. This readout was a one-step, deterministic center-out probe to eight targets at a radius of 0.75, which began from the state the agent would carry into its next reach.

### 4.2 Object-location and object-recognition memory simulations

#### Environment and observation model

The agent moves in a circular arena of radius 1.0 containing two objects. Its state comprises its own heading, a bearing and relative size for each object, three shape features per object (curvature, elongation and angularity), and proprioceptive flags for contact and wall proximity. It never has access to its own position. Object information is egocentric, each object given by its bearing relative to the agent’s heading. The action that the agent takes is a speed and turn rate. The agent follows a smooth random walk whose policy never references object positions, so trajectories are identical in distribution across tasks, conditions and perturbations. Its step is 0.04 arena radii, crossing the arena several times within a session, and its turn rate carries momentum across steps so the path curves rather than jitters.

The position of the object is read from the bearing and apparent size; the identity of the object is read from the shape vector by whisker contact rather than vision. Each sensory channel is degraded in proportion to how well it can currently be sensed,

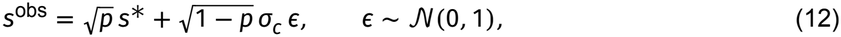

where σ_*c*_ is that channel’s standard deviation. The precision *p* is exact within the sensor’s range and falls off beyond it, bounded below, so a reading taken far from an object leaves a noisier trace than one taken close by. Bearing additionally fades as an object moves away from the agent’s heading, read cleanly to about ±110^°^ and degrading progressively beyond. Visual radius was 0.90 and whisker reach 0.80 in arena units. Heading, contact and wall proximity are never degraded, and nothing is ever substituted: a channel the agent cannot sense becomes noisy. These precise values are not critical: we repeated the full protocol across a sweep over the angular extent, the visual radius and the whisker reach and the dissociation holds throughout (Extended Data Fig. 9).

#### Task protocol

Each trial comprises two 200 step sessions. Either one object is displaced (OLM) or replaced with a new object (ORM); objects are never identical, and the displaced object is moved to a position not previously occupied. MTL’s recurrent state and the cerebellar prediction are carried across the break without reset. All reported quantities are means over 10 independently trained networks.

#### Training

Networks were trained end to end to minimize the cerebellar next state prediction error with Adam, batch size 64, sequences of 200 steps, 60 epochs of 100 updates, learning rate 10^−3^ on a cosine schedule to 10^−5^. On 5% of training sequences, at a step drawn uniformly from 40 to 160, the object was altered partway: displaced on half of those sequences and replaced with a new object in the other half, so that the change is not entirely novel at the test and so both tasks receive identical prior exposure to change.

#### Perturbations

We apply isotropic Gaussian noise of standard deviation σ in three step bursts, once every sixty steps to match the transient perturbation protocol of the Zeidler et al. ^14^ study. Three sites were tested — the cerebellar output, the cerebellar error channel, and the MTL recurrent state — on the same schedule. σ is expressed as a fraction of the perturbed quantity’s own standard deviation, so it means the same thing at each site; the MTL injection is additionally divided by 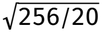 so that the 256-dimensional recurrent state and the 20-dimensional state vector receive the same total injected variance. Figures 7g,h sweep σ from 0 to 0.8 at each site and Extended Data Fig. 8 extends it to 5.

#### Preference index

Prediction error was accumulated separately over the sensory channels belonging to each object and averaged over the *τ* steps following the change, giving *e*_changed_ and *e*_unchanged_. The preference index is

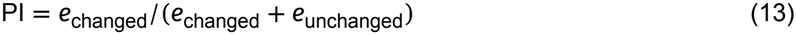

which equals 0.5 when the change leaves the error unaffected. We kept *τ* = 4 throughout except in (Extended Data Fig. 8), where we test timesteps needed to see an effect. Preference retained is (PI_σ_ − 0.5)/(PI_intact_ − 0.5).

### 4.7 Place Cell Experiments

#### Environment and observation model

The agent forages in a circular arena of radius 1.0 with a cue along the wall, matching Rochefort et al. ^13^. The agent’s state comprises its heading, cue bearing with respect to the agent, cue’s intensity and contact flags for the wall and for the cue. It never has access to its own position. Actions and the movement policy are same as used above for Section 4.6. The arena is circular and therefore has no intrinsic orientation, which makes the cue the only allocentric reference. Each sensed channel is degraded with noise using the same rule as described in the OLM-ORM simulations (Section 4.6).

#### State input during darkness

To simulate the unlit condition, which included light extinction and cue removal, we set the sensing radius to zero, so it cannot be touched. In the dark the agent can no longer see the cue, and it can no longer read its own heading. It must therefore fall back on what the cerebellum predicts, and we substitute the five visual channels — the three cue channels and the two heading channels — with the cerebellum’s own prediction of them. Heading in the dark is thus not lost but dead reckoned, as head direction signals are maintained by integrating self motion when visual landmarks are unavailable. The substitution is applied identically to both genotypes. The two contact channels are never substituted because the whiskers work in the dark.

#### Task protocol

Testing follows the five session design of Rochefort et al. ^13^: two cued sessions, two probe sessions, and a final cued session, a probe being either cue present or cue absent. The agent is removed and reinserted before sessions 2 and 5 only. Reinsertion jitters position by 0.05 arena radii and heading by 0.20 rad. Each session is 100 steps and comprises 256 trials. All reported quantities are means over 10 independently trained networks per genotype.

#### Training

Training follows Section 4.6. Each episode additionally contains one blackout of 30–80 steps, roughly 30% of the episode, so that darkness is a regime the forward model is trained for rather than a shift away from its training distribution. The sensing radius was varied across episodes.

#### Genotypes

Wild type networks transmit the cerebellar prediction intact. The L7-PKCI mutant is modeled by corrupting what the cerebellum transmits with noise of standard deviation 0.30 in units of the signal, resampled per sequence. Unlike the OLM-ORM simulations, where Zeidler et al. ^14^ stimulate acutely only during test, L7-PKCI is a genetic manipulation present throughout training — to match this, we apply the cerebellar perturbation during training, so the mutant network learns with a cerebellum that was never reliable. The noise is applied to the cerebellar state prediction.

#### Analysis

Rate maps were built by dividing the arena into 3 radial by 8 angular bins measured from the cue, and averaging each unit’s activity over the times the agent visited each bin; bins visited fewer than four times were discarded. For the example maps, units were ranked by the spatial variability of their map in a cued session and the most variable were taken and plotted across the five sessions so that the same units are shown throughout; each map is scaled to its own peak. Three measures follow Rochefort et al. ^13^. Mean field rate is a unit’s average activity inside its field, the field taken as the bins exceeding that unit’s own map mean by one standard deviation. Spatial coherence is the correlation between each bin and the average of its eight neighbors, which is high when a field is a single contiguous patch. Between session stability is the correlation between two sessions’ maps for the same unit at matched locations. The first two measure how sharp a field is and stability requires the map to stay stable, so a map that drifts coherently scores low even when its fields remain sharp.

We also compared each unit’s dark map directly against its light map using four similarity measures: uncentered cosine, pearson correlation, cosine after rectifying at the light map’s mean, and population-vector correlation across units within each bin. Unlike the measures above these need no per-unit field definition, so they test whether the genotype difference depends on how a field is defined.

### 4.8 Serial reversal task

#### Task

On each trial one of the four stimuli is presented and the agent selects one of the two operants, hold or release. Which operant is correct, and whether performing it correctly pays, is decided by the pair (stimulus, context). There are two contexts and the switch between them is never cued. The outcome is +1 when the correct operant is made on a rewarded stimulus, 0 when it is made on an unrewarded one, and −1 whenever the operant is wrong, matching the time out design in the Bernardi et al. ^8^ task. We use the mapping of Bernardi et al. ^8^: at a switch the correct operant reverses for all four stimuli while the reinforcement contingency reverses for two, so context, operant and reinforcement are mutually orthogonal across the eight (context, stimulus) conditions and no single variable identifies the context. Stimuli are drawn independently and uniformly on every trial. Each session is 240 trials, and the context switches every 50–70 trials, drawn uniformly.

#### State, action and prediction error

The state is the stimulus, one hot in four dimensions; the action is the operant. The cerebellum is a forward model of the outcome, 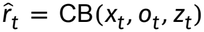, a single scalar. The prediction error is

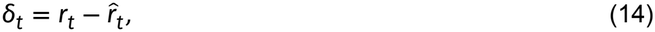

a reward prediction error rather than a state prediction error. MTL receives [*x*_*t*−1_; *o*_*t*−1_; *δ*_*t*_]: it is one step behind on everything except the error, exactly as in the circuit described in sections above.

#### Training

MTL and CB were trained jointly to minimize the mean squared reward prediction error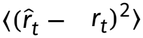, Operants were drawn as coin flips during training, giving an unbiased coverage of both operants at every stimulus. Each circuit was trained for 60 epochs on 2000 sequences of 240 trials, in mini-batches of 128, with gradients computed by backpropagation through time over the full sequence, using Adam at a learning rate of 10^−3^. The prediction error was monitored on 300 held-out sequences. Ten circuits were trained per arm, differing only in seed, and all reported quantities are means over those ten. Two arms were trained. In the *main* arm MTL’s only route to the outcome is the prediction error. In the *augmented control* arm the previous outcome *r*_*t*−1_ is given to MTL directly, so the error is redundant.

#### Perturbations

Weights were frozen and the prediction error reaching MTL was manipulated at test only. In the *zero* condition *δ*_*t*_ was replaced by zero, which removes the cerebellar prediction error entering MTL. In the *shuffle* condition, *δ*_*t*_ was replaced by the error from a randomly chosen other sequence at the same trial and tests whether MTL needs the correct error or merely an error input from any trial.

#### Context decoding and Representational geometry

The eight (context, stimulus) conditions were analyzed with the measures of Bernardi et al. ^8^, on trials at least five from a switch, with up to 400 trials per condition. Decoding is four-fold cross-validated logistic regression on a balanced split of the eight conditions. CCGP trains the decoder on three conditions from each side of a split and tests it on the held-out ones, averaged over all such choices; it is high only when the split generalizes to conditions the decoder never saw. The parallelism score is the mean cosine between coding directions, maximized over pairings of the two sides. Shattering dimensionality is the mean decoding accuracy over all 35 balanced four-versusfour splits of the eight conditions. Three of those splits are named: context, value and action.

Measures were computed for the recurrent state of MTL and for both hidden cerebellar layers. The MTL’s is scored against the variables of the previous trial and the cerebellum against the current trial’s, because the state of MTL is built from [*x*_*t*−1_; *o*_*t*−1_; *δ*_*t*_] and has not seen the current stimulus, whereas the current stimulus and the operant are the inputs of the cerebellum.

## Acknowledgements

We thank members of the Gershman lab for their helpful comments on framing and figures.

## Author contributions

E.A.A. conceived the project and developed the model. C.B. and E.A.A. designed the analyses, ran simulations, and prepared the figures. S.J.G. and E.A.A. supervised the project and acquired funding. All authors contributed to writing and editing the manuscript.

## Code availability

. Simulation and analysis code is available at https://github.com/cheshta2205/motor-and-structure-learning-.

## Extended Data

**Extended Data Figure 1.**
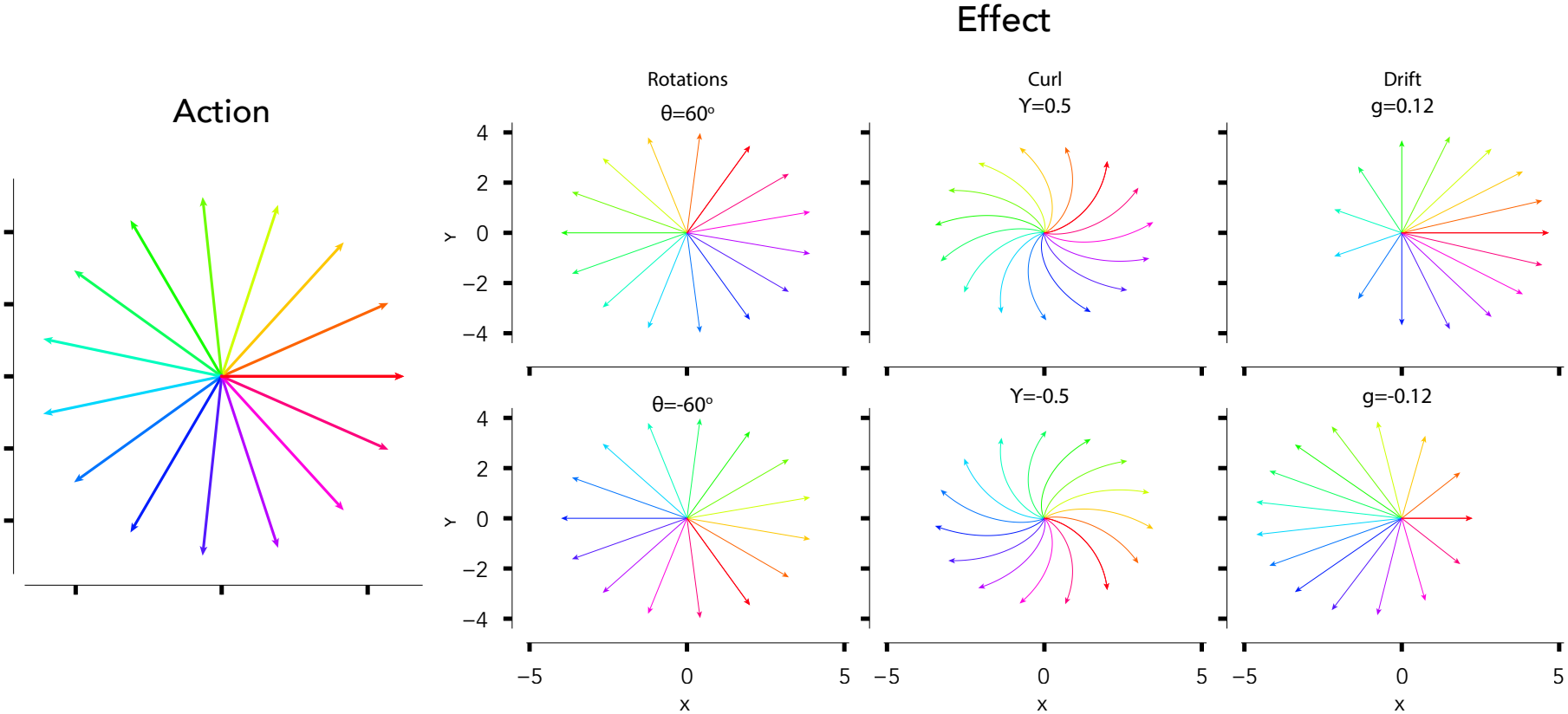
Context-dependent effects of action. **a**, Twelve action directions available to the agent, each a constant force held for sixty steps. **b**, The resulting trajectories under each family of perturbation at two values of its latent parameter: rotations of ±60°, curls of ±0.5, and drifts of ±0.12 per step. The same action produces a different outcome in each context, so the mapping from action to consequence cannot be learned without inferring which context is active.

**Extended Data Figure 2.**
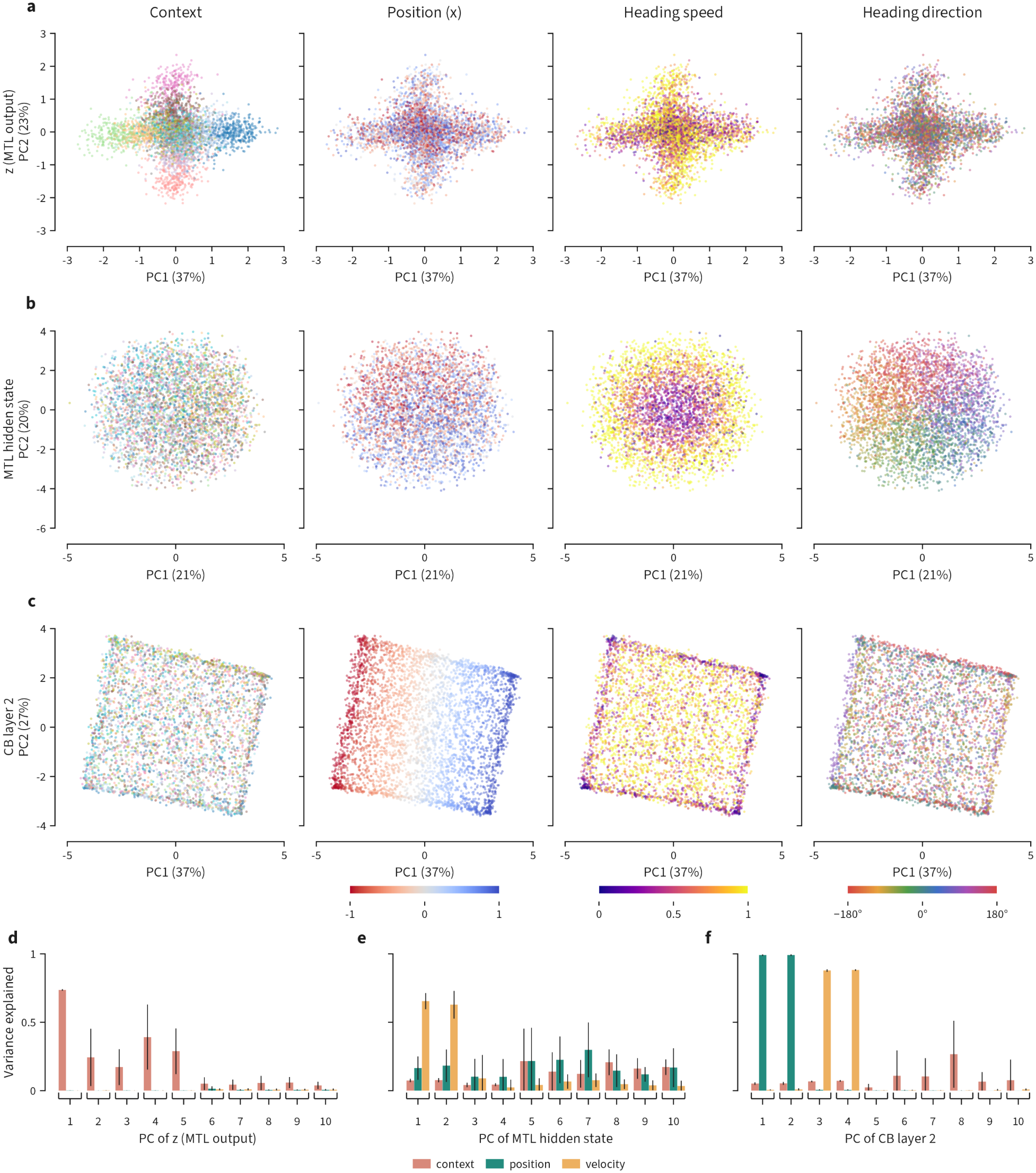
Representational analysis of the CB-MTL circuit. **a**, The MTL latent in its top two principal components, colored by context, position, heading speed and heading direction. Experiences cluster by context. **b**, The same projections for the MTL hidden state, which instead reflects the agent’s heading speed and direction. **c**, The same for the second cerebellar layer, which reflects the agent’s position. **d–f**, Proportion of each principal component’s variance explained by context, position and velocity, for the latent, the MTL hidden state and cerebellar layer 2 respectively. Context accounts for over half the variance of the leading components of the latent and for almost none in the other two layers, where position and velocity dominate.

**Extended Data Figure 3.**
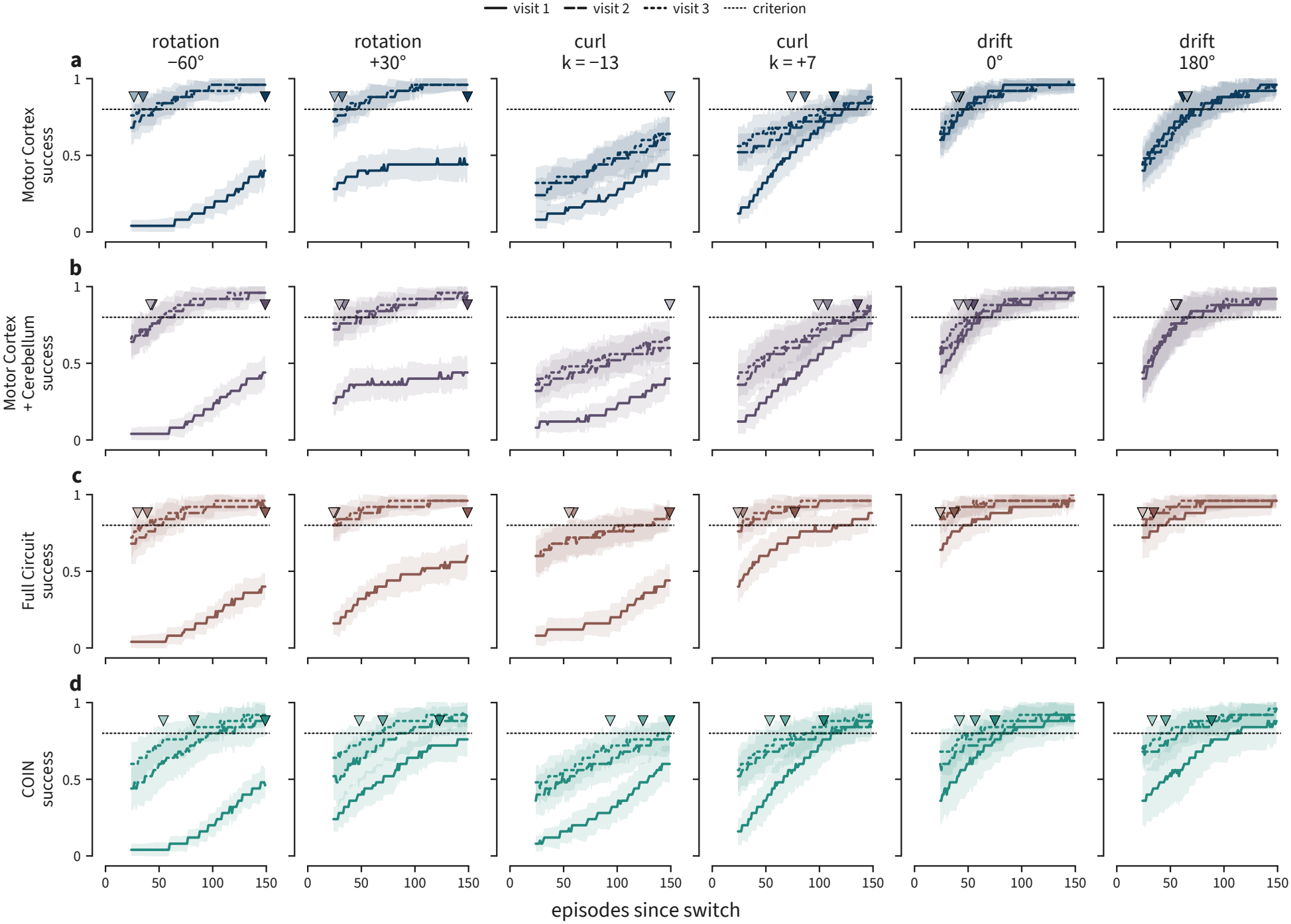
Per-context learning curves across three visits. Success against episodes since switch for each of the six training contexts, shown separately for each circuit: **a**, MC. **b**, MC + CB. **c**, CB–MTL. **d**, COIN. Solid, dashed and dotted lines give the first, second and third visit; the horizontal dotted line marks the 80% criterion and triangles mark the episode at which each visit reaches it. Lines denote median across seeds, and bands are the interquartile range across seeds.

**Extended Data Figure 4.**
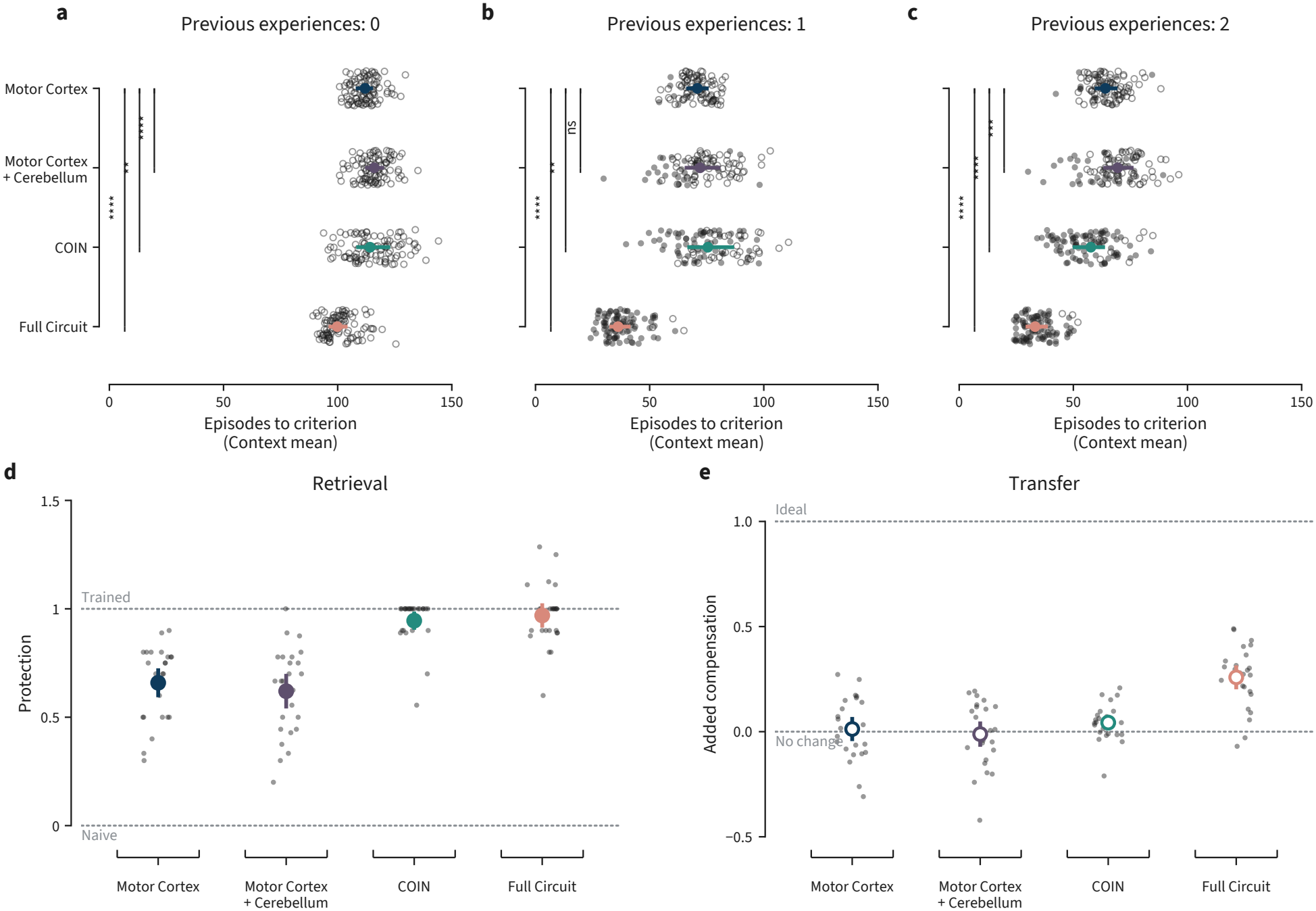
CB-MTL circuit has high transfer and high retrieval. **a–c**, Episodes to criterion in the six-context cycle, by the number of previous exposures to the context. Criterion is an 80% success rate, evaluated on the 25-episode running mean. One point per seed, giving its mean over the six context blocks; point and whisker, median and interquartile range over seeds. Seeds whose means contain blocks that never reach criterion are drawn open. Brackets are two-sided Mann–Whitney *U* tests against MC, unpaired, one value per seed, Holm-corrected within each panel. **d**, Retrieval, measured as protection from interference: (revisit−naive)/(trained−naive) in success over the first ten episodes, so that 1 is survival of the trained solution through a conflicting rotation and a washout and 0 is a return to naive. **e**, Transfer, measured as the compensation added over the first ten reaches: (aim − parked aim)/ideal aim, so that 1 is full compensation beyond where the policy started and 0 no change. Both from the rotation pair, −60^°^ then +30^°^ trained, followed by a washout; median and interquartile range over seeds. Two-sided Mann–Whitney *U*, unpaired, Holm-corrected within each panel: in **d**, each context-inferring model against each context-blind one; in **e**, the CB–MTL circuit against each other model. All *p* < 10^−4^.

**Extended Data Figure 5.**
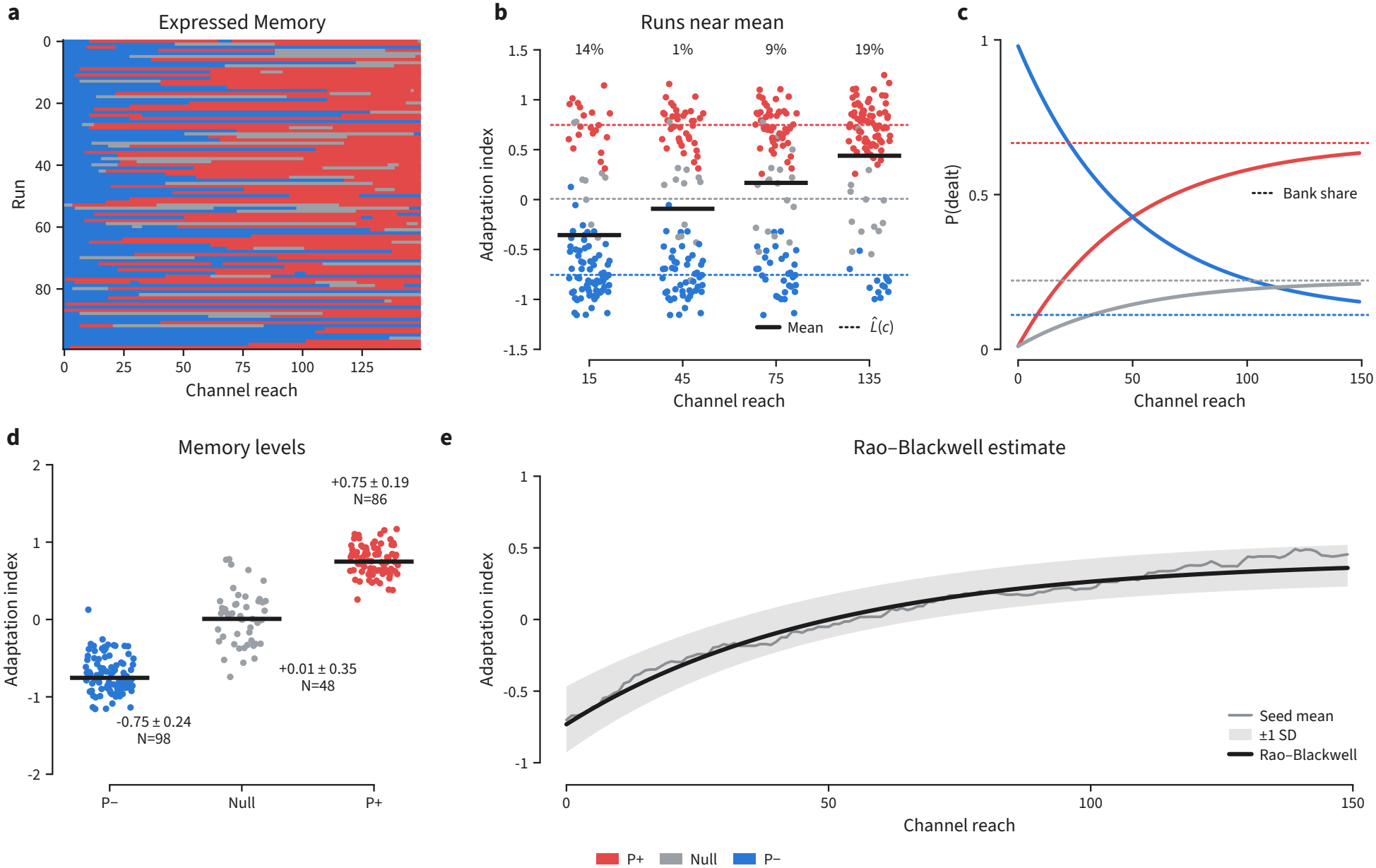
Spontaneous recovery belief decomposition. **a**, Which memory is expressed on each channel reach, one row per run. Null (gray) corresponds to the neutral reach condition under no visuomotor rotation while P+ (red) and P-(blue) correspond to the 60° and −60° rotations. At each reach the MTL state is re-initialised from a uniform draw over the buffer with probability *h* = 0.02, and otherwise carries over from the previous reach. **b**, Adaptation index at four channel reaches, *k* = 15, 45, 75, 135; dotted lines give 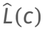 the Rao–Blackwellised adaptation index conditioned on each memory, and solid bars the mean. Percentages give the fraction of runs within ±0.15 of that mean. **c**, Probability that a given memory is the one expressed. The weight on the memory carried over from the start of the channel block decays as (1 − *h*)^*k*^ with *h* = 0.02; the remaining weight rises toward the bank share, the composition of the bank of past MTL end-states, in which older states are down-weighted by a power law. **d**, 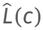 for each of the three memories, with the number of runs in which each was expressed; a run may be dealt more than one memory over the block. **e**, Mean adaptation index across the channel block, 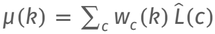 compared against the sampled realization and its spread across 100 runs. The recovery is the deal weights moving from the carried-over *p*− state toward the buffer’s composition, not a change in any 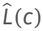.

**Extended Data Figure 6.**
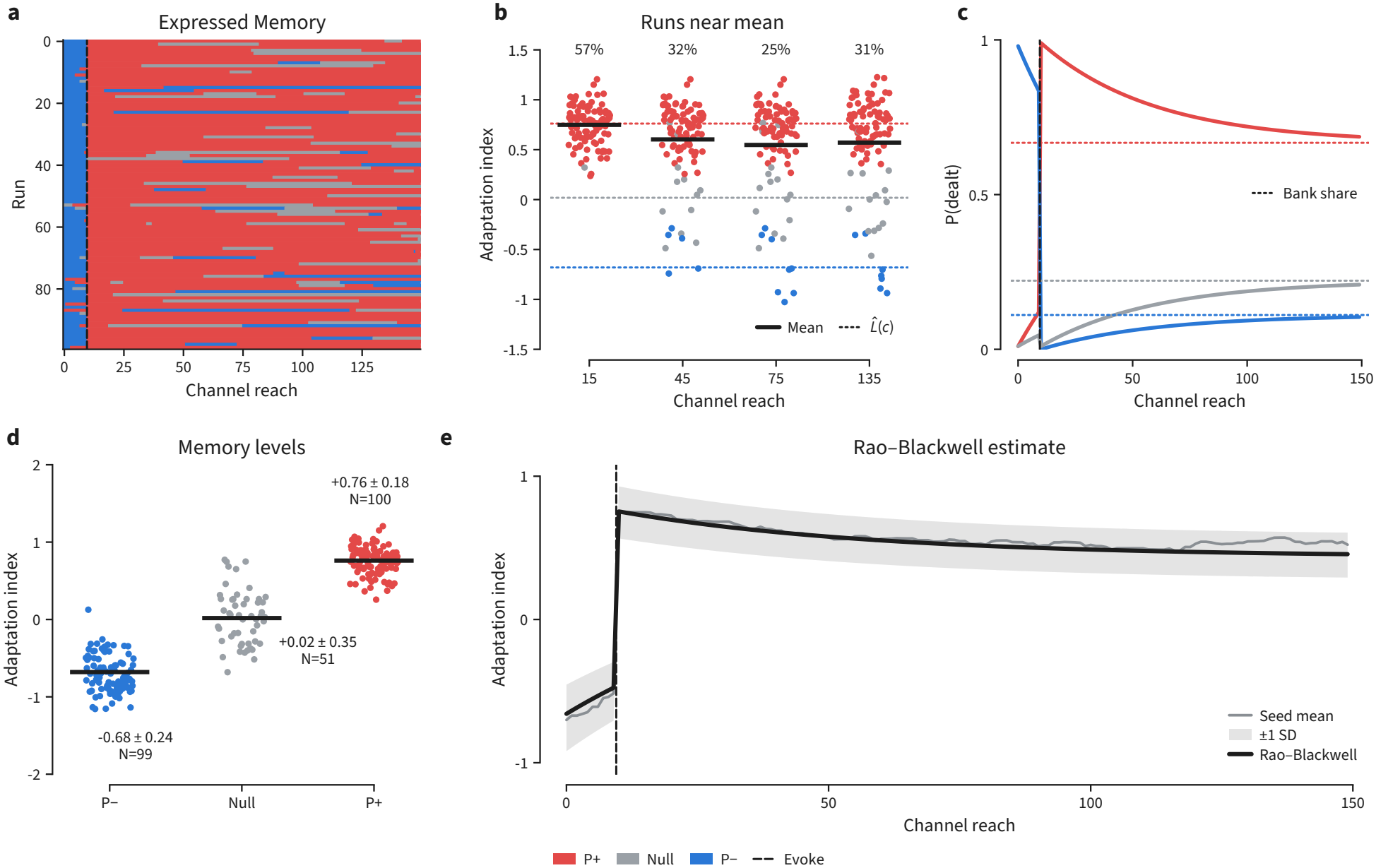
Evoked recovery belief decomposition. As in Extended Data Fig. 5, for the evoked recovery paradigm; the dashed line marks the evoking reach. **a**, Memory expressed on each channel reach, one row per run. **b**, Adaptation index at *k* = 15, 45, 75, 135, with 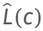 and the fraction of runs within ±0.15 of the mean. **c**, Deal weights; the single *p*+ exposure resets the carried-over state to *p*+, so the weight on *p*+ starts near one and decays toward the bank share rather than rising toward it. **d**, 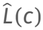 per memory with run counts. **e**, *µ*(*k*) against the sampled realization. The jump and subsequent decay are the same deal process run from the opposite initial condition.

**Extended Data Figure 7.**
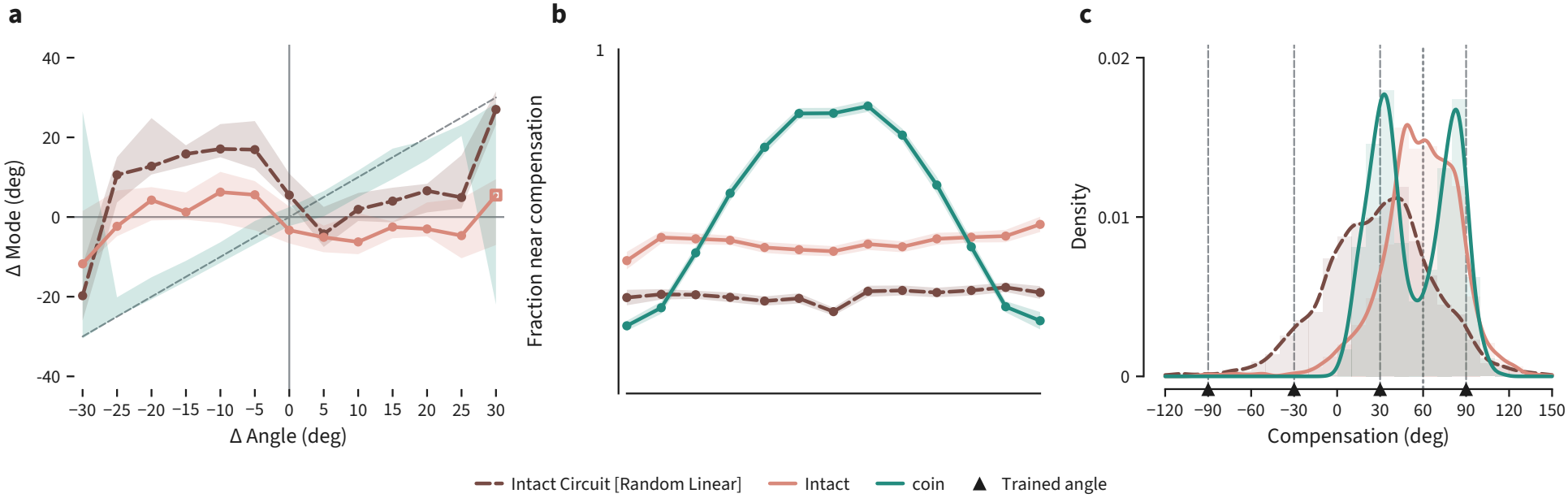
Single-trial compensation distinguishes discrete and continuous representations. All panels use the frozen sweep of section 2.6, in which no weights are updated. **a**, The peak of the per-reach aim distribution at each rotation, as an error from full compensation, against that rotation’s signed offset from the nearest trained anchor. A readout that commits to the nearest anchor on each reach falls on the identity line, while a continuous axis stays near zero. **b**, Fraction of reaches aimed within 15° of full compensation, against the same offset. **c**, Distribution of per-reach compensation at ±60°, the midpoint between two anchors, folded so that full compensation reads 60°; triangles mark the trained angles. COIN’s distribution is bimodal, with modes at the two nearest anchors, while both circuit variants are unimodal near the true angle. Mean compensation at this rotation is nearly equal for the circuit and COIN, so the difference is in the single-trial aim rather than in average performance.

**Extended Data Figure 8.**
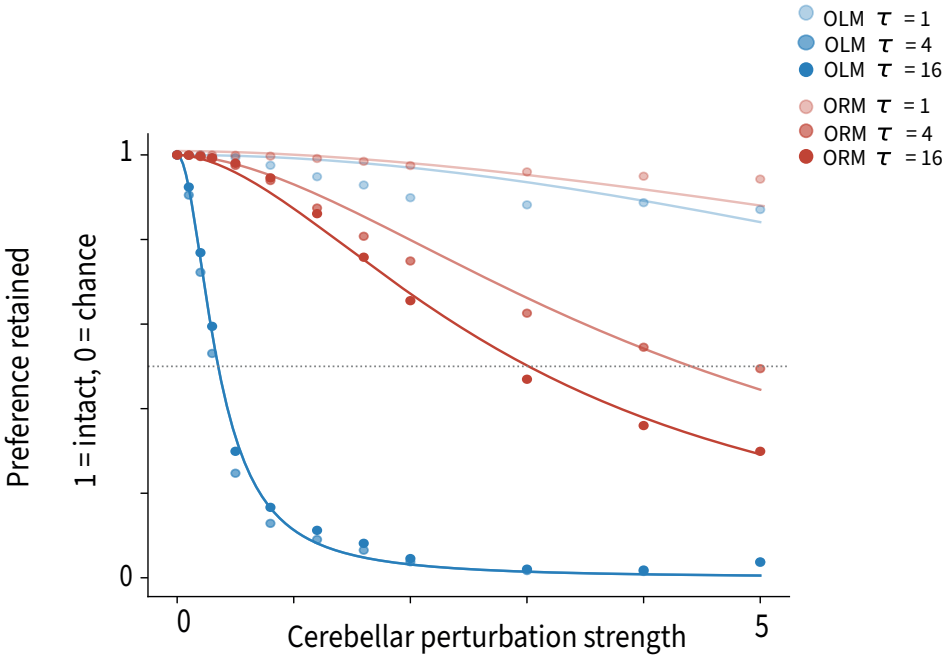
The dissociation does not depend on perturbation strength or averaging window. Preference retained as a function of cerebellar perturbation strength σ_CB_, for OLM (blue) and ORM (red), at three post-change averaging windows (*τ* = 1, 4 and 16 steps; light to dark). The retained preference is the normalised preference index within each task and window, so that 1 is the intact value and 0 is chance.

**Extended Data Figure 9.**
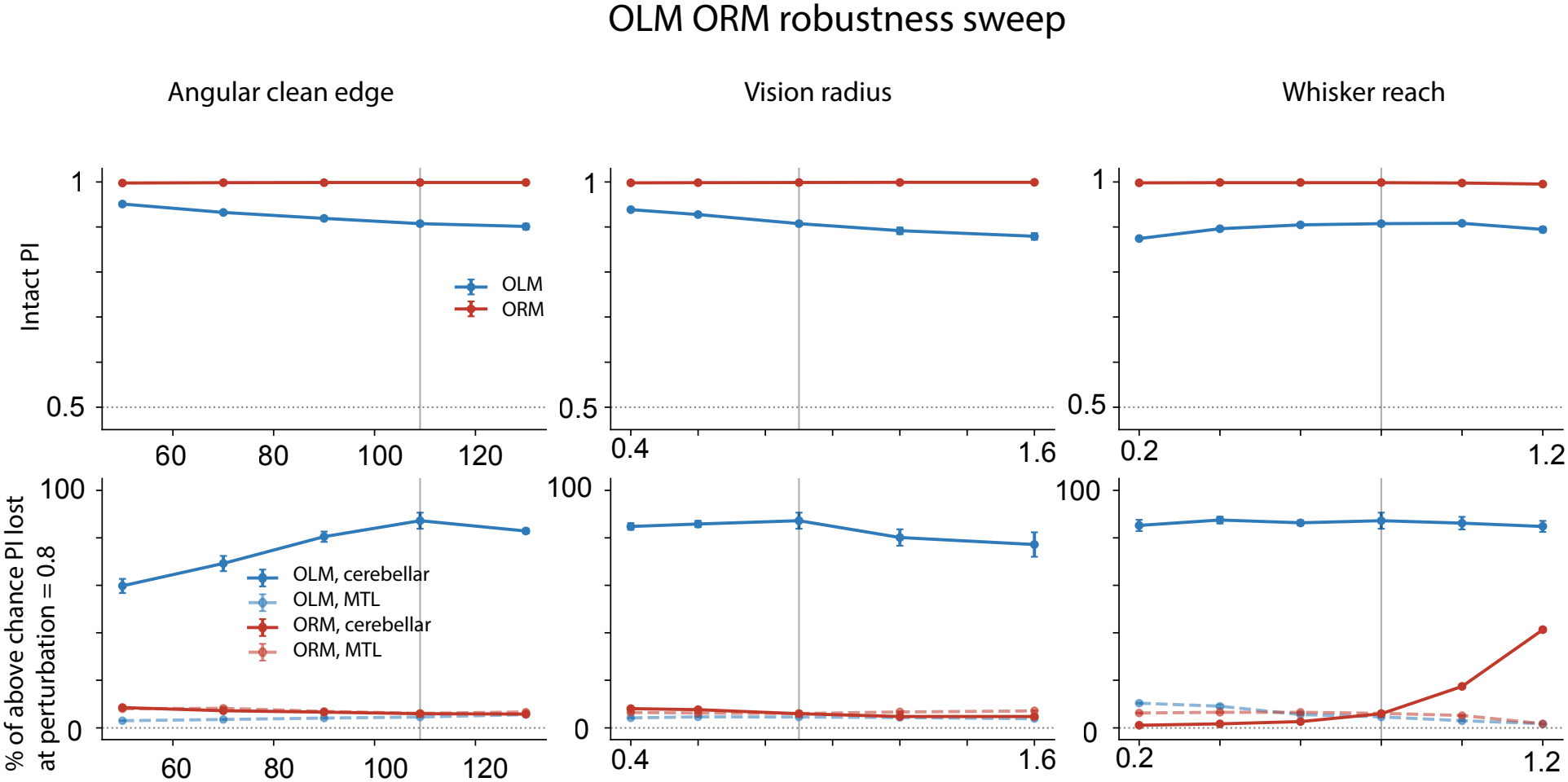
Robustness of the OLM/ORM dissociation to the observation model. **(a–c)** Intact preference index against the angular extent over which bearing is read cleanly, the vision radius, and the whisker reach within which shape is sensed exactly. **(d–f)** Percentage of the above-chance preference index lost at σ = 0.8, for cerebellar and MTL perturbation in each task. Vertical lines mark the setting used throughout the paper. Each point is the mean of three independently trained networks; error bars give s.e.m.

**Extended Data Figure 10.**
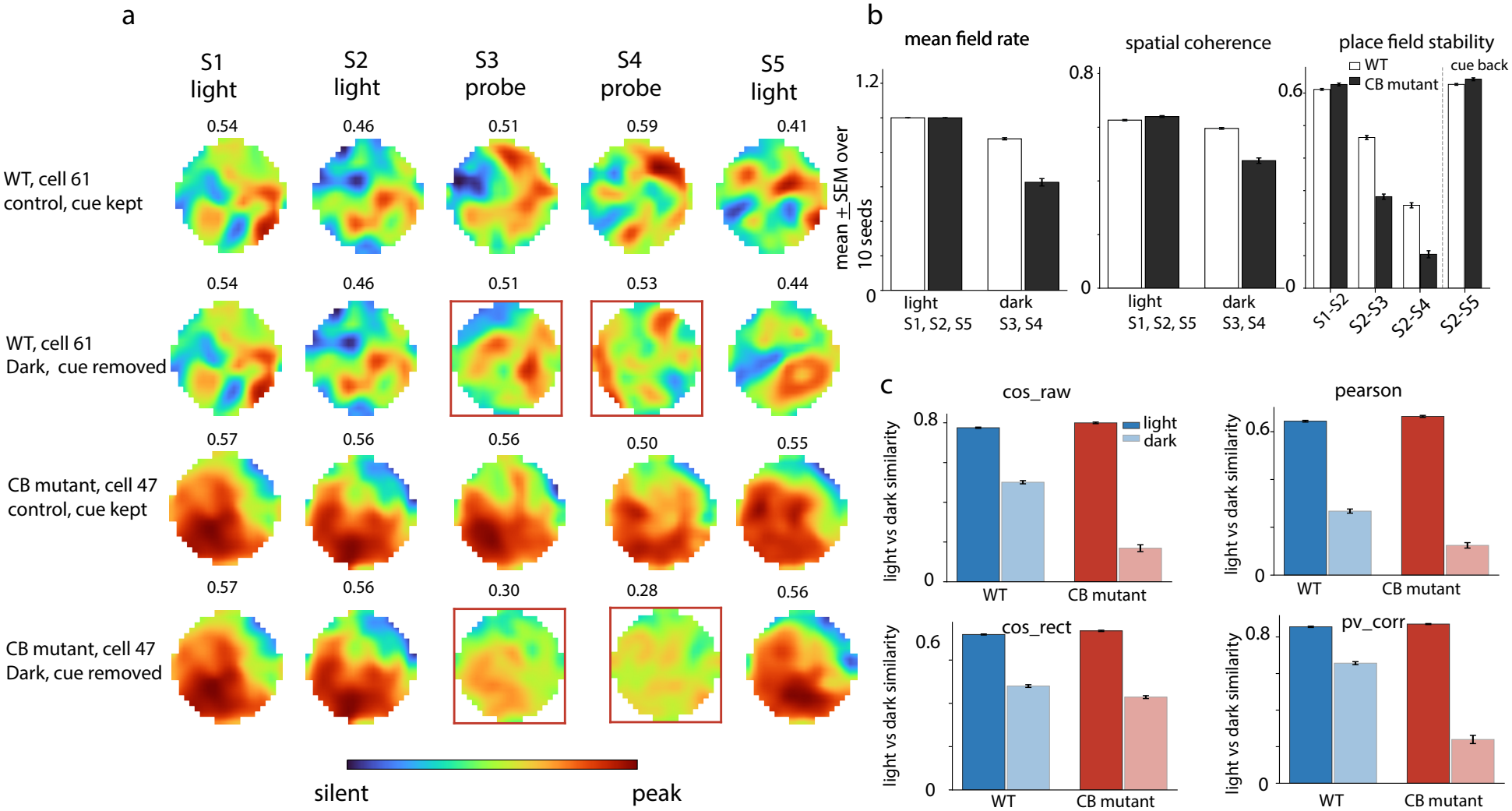
Place field stability under cerebellar perturbation. **(a)** Rate maps for one cell across the five-session protocol (S1, S2 and S5 cued; S3 and S4 probe), shown for wild type and CB mutant circuits with the cue kept in S1-S2 and with it removed in S3–S4, and returned in S5. Red boxes mark the probe sessions; numbers give peak rate. **(b)** Mean field rate and spatial coherence in cued and probe sessions, and between-session place-field stability, including recovery once the cue is restored. **(c)** Similarity between cued and probe maps on four measures: raw cosine, Pearson correlation, rectified cosine and population-vector correlation. Mean ± s.e.m. over ten networks.

**Extended Data Figure 11.**
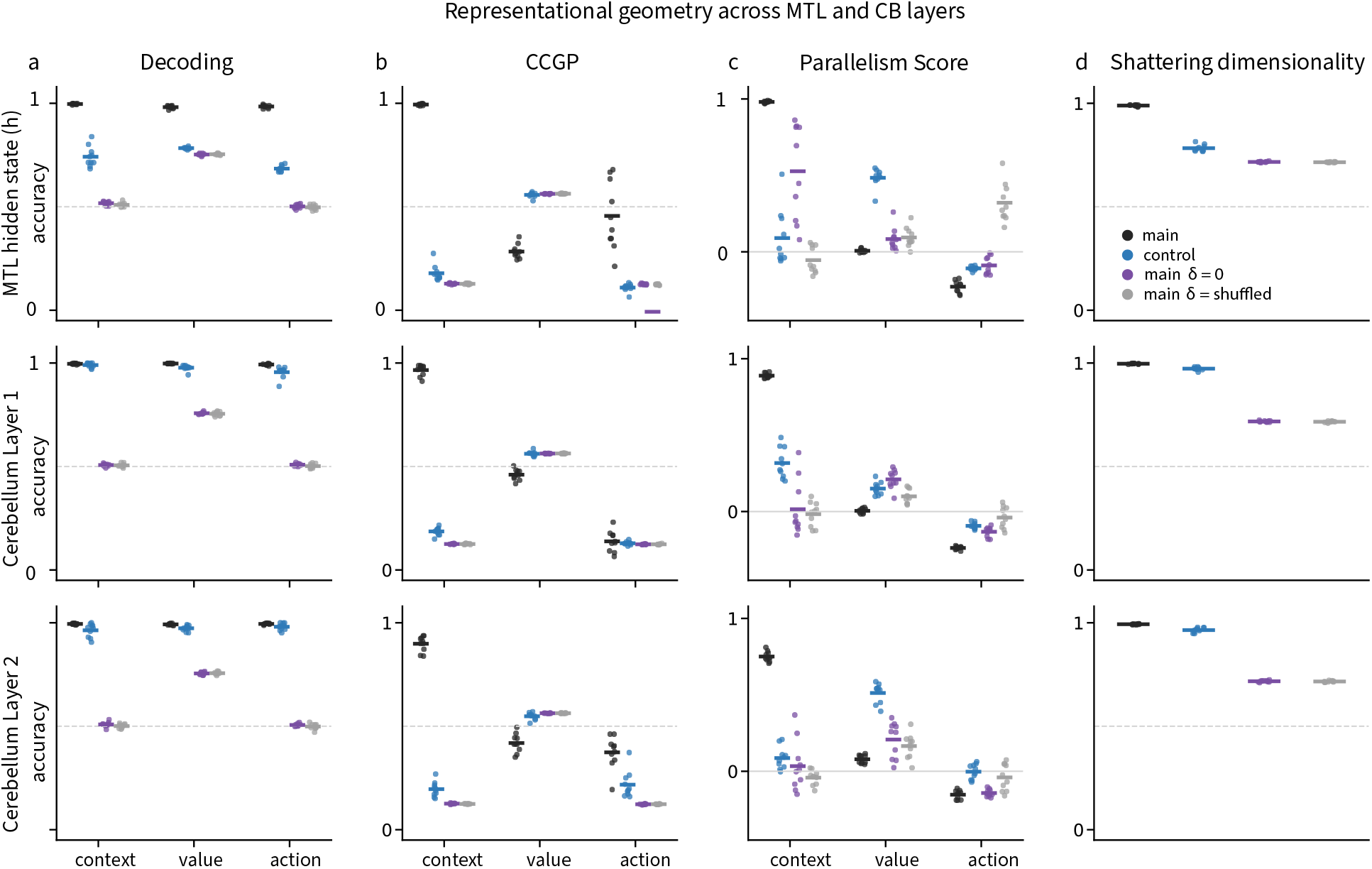
Representational geometry across MTL and cerebellar layers for the SRO task. Decoding **(a)**, cross-condition generalization **(b)**, parallelism score **(c)** and shattering dimensionality **(d)**, computed for the MTL hidden state and for cerebellar layers 1 and 2. Four conditions are shown: the main arm, the augmented control arm that additionally receives the previous reward, and the main arm with the cerebellar reward prediction error replaced by zero or shuffled across sequences. Points are individual seeds; horizontal bars give the mean over ten networks. Dashed lines mark chance.

